# Rapid, behaviorally timed increases in blood-brain barrier permeability are predicted by endothelial calcium events and driven by histaminergic axons

**DOI:** 10.64898/2026.11.10.765662

**Authors:** Kevin L. Turner, Seneca J. Scott, Sinda Fekir, Eric Teboul, Kat Browning, Chanel I. Johnson, Akshay Nagar, Christopher I. Moore

## Abstract

Survival requires a detailed understanding of body state, and the vasculature provides a continuous conduit of this information to the brain. Yet the blood-brain barrier (BBB) is canonically viewed as a stable boundary, and any passage across it as pathological leak, because sustained exposure carries real risk. How the forebrain resolves these competing needs for information versus safety remains unknown. Here we show that the healthy barrier opens briefly and locally at moments when body-state information may be worth the risk. Using two-photon imaging of an intravascular dextran in the somatosensory neocortex of mice performing a cued reward task, we identified ‘Plume Events’: transient, focal increases in permeability at penetrating arterioles that were locked to the cue and reward, persisted on trials without detectable dilation, and shifted earlier with training. Endothelial calcium events preceded and predicted Plume Events within seconds and within 15 micrometers of the vessel wall. Histaminergic axons from the tuberomammillary nucleus were active before Plume Events, and their optogenetic stimulation drove Plume Events that waned across repeated trains while dilation persisted. Together, these findings reframe the blood-brain barrier as a dynamic interface whose openings are immediate, local and transactional, granting the forebrain access to body-state signals at behaviorally relevant moments. Such events may also support brain health through locally timed metabolic supply and waste clearance.

## Introduction

Taking correct actions, and learning from them, requires understanding body state. A thirsty mouse choosing to go, or not go, from its nest to a creek at dusk must weigh dehydration risk against competing drives (e.g., protection of young), and decide if external sensory context (e.g., lack of predator sounds) favors resolution of its needs. After choosing to go, persisting to its goal through evolving context (e.g., a hawk screech when almost there) also requires touching base internally (a ‘gut check’).

Optimal learning similarly requires a high-resolution snapshot of body state. Future success depends on the detail with which learning circuits link an outcome, in a given sensory environment and following a given action, with the many internal variables that describe ongoing needs. When that internal state next emerges and matches the external sensory environment, learning can bias the re-emergence of those ensembles that created correct actions. In sum, choosing, persisting and learning all require a precise snapshot of body state.

The forebrain provides multi-level computation to support such real-world problem solving. Neocortex and striatum integrate internal and external input with a lifetime of prior learning to create rapid, state-appropriate choices and learn from them, i.e., whether a pattern of specific internal needs was satisfied by an action. Vascular networks provide a continuously updating conduit of body state, an ideal ‘highway’ to the forebrain. Metabolic signals^1–3^, hormonal information^4–6^ and immune status^7–9^ are all transmitted in vascular channels, and the dense vasculature throughout the forebrain places it in close proximity to this reservoir. Lower structures such as vagal inputs and the circumventricular organs are chronically permeable to these body signals and respond by reformatting them into action potentials sent to the forebrain and/or hormones delivered back into vessels^10–13^.

However, the forebrain blood-brain barrier (BBB) is believed to prevent rapid access, on the time scale of decision making and trial-based learning, to almost all molecules > 500 Da^14,15^. This blockade is thought essential to brain safety^16–19^, as chronic failures of the BBB are a hallmark of many diseases, including Alzheimer’s disease (AD)^20–23^. As such, transmission of high-dimensional state information through the direct channel of the vasculature is often viewed only as a slow, diffusive regulator of behavior, with the sole ‘fast’ source of interoceptive signals the transmission of action potentials through a cascade of neural areas that ultimately impact the forebrain^24,25^.

We propose an updated view, that the forebrain BBB is dynamic, providing brief, timely access to detailed state information, but only when behavioral value makes it worth the risk to brain health. Discrete moments of permeability in engaged circuits could minimize overall exposure risk, while still delivering real-time signals to recruit the right ensembles for optimal actions, and support learning that links a current state to such ensemble recruitment in the future. We refer to this prediction as the ‘State Gate Hypothesis,’ one of several kinds of vascular dynamics that can play an active role in creating adaptive behavior^26^. In addition to potential behavioral benefit, rapid, local, activity-dependent BBB permeability could also have distinct health advantages. While ‘functional hyperemia’ created by vasodilation is not thought to support metabolic delivery^27^, highly local, tuned BBB permeability—which our findings show can be decoupled from vasodilation—could provide ‘metabolic microsupply’ to engaged representations (e.g., a neocortical mini-column). Transient BBB permeability could also provide rapid, focal waste clearance^28^ during behaviorally relevant processing, helping prevent activity-related toxic buildup. In sum, in support of processes such as information relay, metabolic supply and waste clearance, the State Gate Hypothesis predicts the existence of event-specific relay, on time constants ∼two orders of magnitude faster than those currently accepted.

This prediction is consistent with a more general emerging view that the BBB is adaptive^29,30^, acting as an ‘interface’ more than as a barrier^31–35^. The specific prediction tested here, that the BBB communicates in discrete events on time scales appropriate to ongoing choice and learning, and matched to local need (e.g., information, waste clearance), is supported by decades of studies showing that minutes of continuous sensory drive can increase BBB permeability in sensory neocortex. In 1965, Lorenzo et al.^36^ found that intravenously (IV) injected radiolabel localized to visual thalamus and neocortex after 5 minutes of photic stimuli, and to the auditory system after acoustic drive. Similarly, in anesthetized rats, several minutes of vibrissa stimulation or forepaw shock also increase primary somatosensory neocortex (SI) permeability to interleukins as well as ∼70 kDa Evans blue dye^37^. Recently, these studies using *postmortem* access have been complemented by dynamic contrast imaging studies in humans, using a repeated grasping task for 30 minutes^38^. These activity-dependent assays could reflect the accumulation of many transient events, as described below.

The prediction that BBB dynamics are rapid and discrete diverges from common views: they more closely parallel the discrete, transactional nature of synaptic action potential relay. In this communication mode, rapid voltage shifts (depolarizations) create local axon Ca^2+^ events that release a chemical information signal. Myriad studies show that endothelial cell calcium events (ECCE) can increase BBB permeability in in *vitro/ex vivo* models^39–44^. Paracellular transmission through tight junctions is enhanced by Ca^2+^-mediated endothelial cell contraction^45–47^. Recent discoveries *in vivo* show that ECCE occur at a high rate in SI^48–51^, and, in anesthetized mice, can be driven by sensory input (electric foot shock)^52^.

The production of behaviors critical to survival and associated learning moments most often occurs during high arousal states. Histamine (HA) release from neurons of the tuberomammillary nucleus of the hypothalamus (TMN) is critical to evoking such states^53–56^. In the brain, HA is a transmitter, acting on receptors to influence neural activity. It is synthesized and released by neurons within the TMN, an evolutionarily ancient node within the ascending arousal system^57–60^. The TMN projects brain-wide, releasing HA to support wakefulness, arousal, motivated behavior, and learning and memory^61–63^. The TMN supports invigoration during consumption, and is necessary in driving escape behaviors under threat^64–70^. Indeed, HA has been proposed to act as an ‘alert signal’ in the brain – exactly the moments in which the forebrain would ideally want access to body-state signals^71,72^.

TMN axons are anatomically well-positioned to mediate Plume Events as they directly contact forebrain vessels^73^. HA is also known to rapidly increase BBB permeability through the activation of HA receptors on abluminal (brain-facing) pial vessel endothelial cells^74–83^. These effects are mediated by endothelial cell calcium-dependent signaling mechanisms, which act to rapidly reorganize endothelial cell junction adhesion and increase paracellular permeability^84,85^. Indeed, inhibiting neocortical HA receptors pharmacologically has been shown to tighten the BBB, indicating mechanisms of endogenous regulation of BBB permeability by basal neocortical HA tone^86^. Behaviorally, TMN activity is upregulated during need-state-motivated behavior, and TMN axons are critical to coordinating escape behaviors during threat^65,87^. These are exactly the conditions under which the State Gate Hypothesis predicts the BBB should open. We thus reasoned that rapid, behaviorally tuned HA release from TMN-SI axons might drive Plume Events. We examined this possibility by concurrently recording TMN-SI axon calcium activity and BBB permeability during behavior, and by examining the effects of optogenetic recruitment of TMN-SI axons on local BBB permeability *in vivo*.

Here, we report the discovery of Plume Events. Across task contingencies, Plume Events onset in specific, sub-second behavioral epochs. On the second day of training on a Pavlovian task, Plume Events onset immediately prior to reward (‘pre-reward’), and following reward. With extended training in an instrumental variant of the same task, these distinct Plume Event onset epochs are also observed, along with the emergence of an additional set of Plume Events, immediately following a valid sensory cue. ECCE recorded simultaneously predicted the onset of proximal Plume Events, with larger ECCE GCaMP responses predicting larger effects. We also discovered that TMN axons in somatosensory neocortex rapidly respond to cues and rewards, that optogenetic activation of histaminergic TMN axons drives Plume Events, and that their endogenous spiking predicts them. We discuss this new form of body-to-brain communication in the context of behavior and its potential role in metabolic supply and waste clearance.

## Results

### Plume Events Occur Timed to Key Behavioral Epochs

To systematically test the prediction that the BBB shows rapid dynamics, we employed a well-characterized task in which vibrissa deflections indicate that mice can lick to obtain a water reward (**Fig. 1a**). To track BBB permeability, we conducted two-photon imaging of the movement of IV fluorescent molecules into the neocortex. We employed rhodamine B (RhoB) attached to large dextrans, as these markers are neutrally charged (typically having lower BBB permeability) and lack substantial bioactivity. We chose 70 kDa for these experiments as we found, consistent with prior studies^88^, that the IV availability of 10 and 20 kDa RhoB signal in neocortex declined across tens of minutes, while 70 kDa typically maintained robust signal for > 1 h.

**Figure 1.**
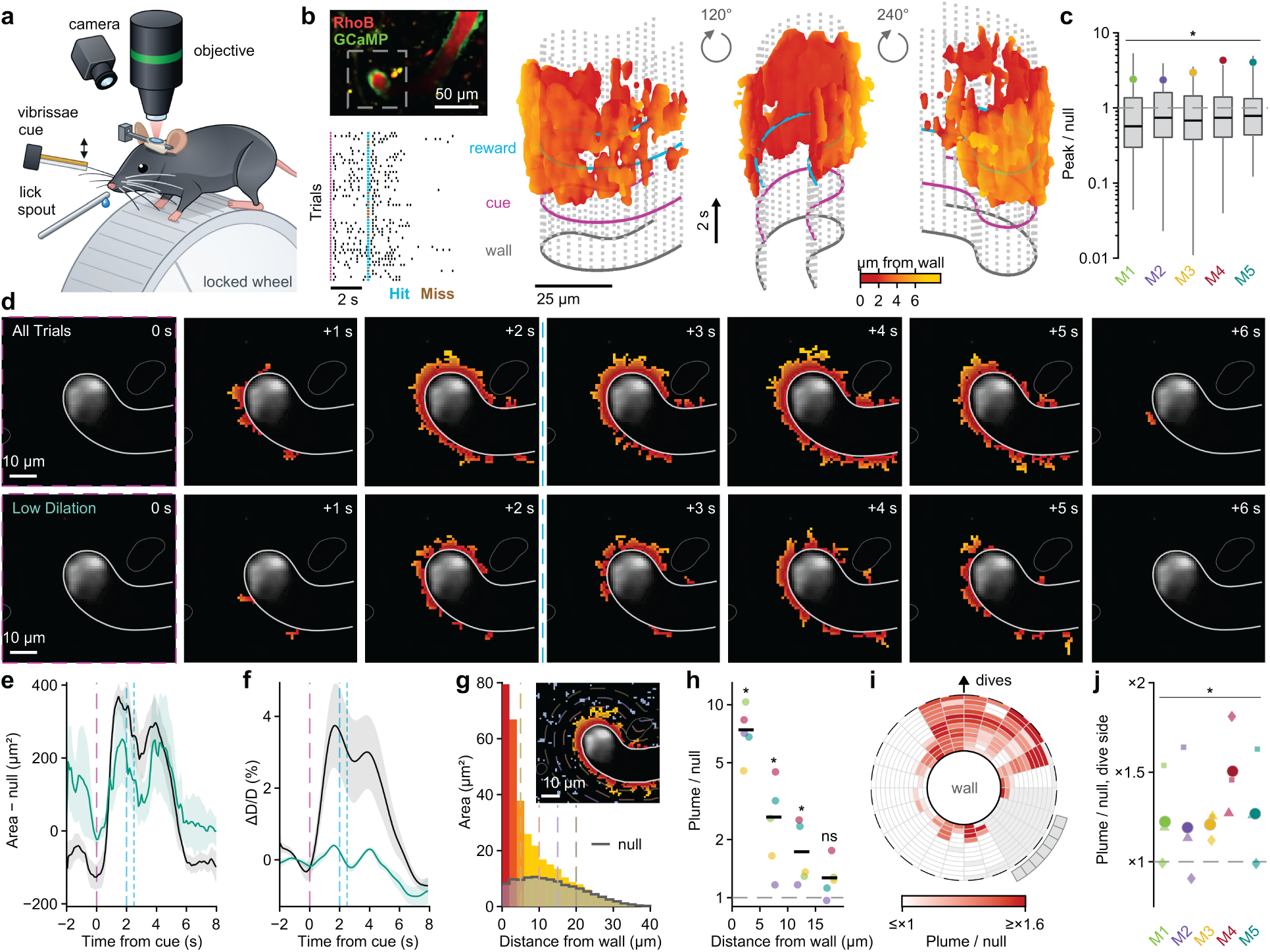
Plume Events are driven by behavior and temporally aligned to relevant epochs a, Schematic of the experimental setup. Head-fixed mice on a locked wheel performed 4 days of Pavlovian (classical) conditioning followed by 8 days of instrumental conditioning during two-photon imaging. Following a vibrissae cue, mice either automatically received the reward (Pavlovian) or were required to lick the spout within 2 s of the cue (instrumental) for the reward to be dispensed. **b**, Inset shows a representative FOV with 70 kDa RhoB-dextran in the blood plasma (red) and endothelial Cdh5-GCaMP8 (green) marking the vessel wall. Below, a lick raster for the first 50 trials of an instrumental session. Cyan and brown ticks mark rewarded (Hit) and unrewarded (Miss) trials. Aligned to this trial structure, a threedimensional reconstruction of the trial-averaged Plume Event from the same session is shown with time on the vertical axis, viewed from three angles 120° apart. Color indicates distance from the vessel wall. Grey dots mark the vessel wall tracked on a frame-by-frame basis, and rings mark the wall at trial start (grey), the cue (magenta) and the reward (cyan). **c**, Peak Plume Event area during the reward window relative to a null distribution built from 100 trial averages of the same session aligned to random times rather than the cue. Circles represent individual mice and boxes show the distribution of random-alignment peaks. Values above 1 indicate a Plume Event larger than expected by chance. **d**, Cue-aligned trial average from the session in b at 1 s intervals following the cue. Top: all trials (n = 164). Bottom: Low Dilation trials only (n = 129), a subset whose mean arteriole diameter stayed within ±0.5% of its pre-cue baseline for 5 s following the cue. Plume pixels are colored by distance from the tracked vessel wall (light grey). The cue frame is outlined in magenta and the dashed cyan line marks the reward. **e**, Plume area following the cue across mice, measured as the area above each session’s random-alignment null. All trials (black) are shown alongside Low Dilation trials (teal; 10 of 15 sessions), selected as in d to separate the Plume Event from arteriole dilation. **f**, Arteriole diameter (ΔD/D) for the same trials as in e. The average of all trials dilates following the cue, while Low Dilation trials stay flat by construction. **g**, Area of pixels above threshold during the reward window as a function of distance from the vessel wall, in 2 µm bins. The grey line shows the same measure for the random-alignment null, which reflects the amount of tissue available at each distance. Inset shows pixels above threshold across the region of interest of the session in b, with dashed lines 5, 10, 15 and 20 µm from the wall. Patches rejected by the Plume Event criteria are shown in blue-grey. **h**, Fraction of pixels above threshold in bands 0–5, 5–10, 10–15 and >15 µm from the vessel wall, relative to the randomalignment null. Values above 1 indicate more Plume Event than expected by chance. Circles represent individual mice and black bars the population average. **i**, Map of Plume Event density relative to the random-alignment null around the vessel wall, averaged across mice. Each vessel was rotated so that the side where it dives out of the imaging plane points up, and mirrored so that its parent segment (grey wedge) lies to the right. White indicates density at or below chance. j, Fraction of the Plume Event within 45° of the dive side relative to the random-alignment null. Small symbols represent individual sessions (triangle, Pavlovian Day 2; square, Instrumental Day 2; diamond, Instrumental Day 8) and large circles the average for each mouse. Values above 1 indicate a bias toward the dive side. n = 5 mice, 15 sessions (Pavlovian Day 2, Instrumental Day 2 and Instrumental Day 8 for each mouse). Colors denote individual mice. Shaded regions indicate ± SEM. Dashed magenta and cyan lines mark the cue and reward. *P < 0.05, ns, not significant; exact one-sided Wilcoxon signed-rank test (c) or sign-flip test (h, j) across mice.

In all fields of view (FOV), we centered imaging on a pial arteriole from its branch point until it sharply descended into the neocortex (the pial-to-penetrating arteriole, PiPA), at the layer I-II border. This common target provided consistency across experiments and includes the Virchow-Robin space, an extravascular region surrounding the penetrating arteriole that has been proposed to be a robust integration zone^89,90^. Further, recent data suggest that this vascular segment may have a more permissive BBB^91^, in part through expression of caveolae that can mediate transcellular transport, not found in brain capillaries^16,92,93^. As such, this position seemed the ideal starting point for testing the rapid, behaviorally timed BBB permeability event hypothesis.

This paradigm revealed ‘Plume Events,’ brief periods of enhanced BBB permeability timed to behaviorally important epochs. In our first study (**Supplementary Fig. 1**), mice were imaged during vibrissa deflections that predicted a water reward 2 s later, on the second day of Pavlovian training that did not require a response for reward delivery. We first analyzed these data by trial-based averaging (∼80 trials/mouse) of fluorescence in a region of interest (ROI) adjacent to the penetrating arteriole. These data showed three distinct increases in fluorescence, immediately after sensory onset, ∼0.5 s before reward, and ∼1 s after reward.

We next imaged a second cohort trained on an instrumental version of the task, in which a response during the post-stimulus window was required to receive reward. To systematically map the onset, origin and trajectory of increased BBB permeability, we employed an unbiased clustering strategy, Density Based Spatial Clustering of Applications with Noise (DBSCAN^94^). This non-parametric algorithm does not require pre-specification of spatiotemporal shape, or pre-definition that any clusters, or a set number, exist (see ***Methods***). When we applied this clustering to the mice from both cohorts (n = 3 and 5), we observed spatially distinct Plume Events that onset ∼1.5 s after sensory stimulation and 0.5 s before reward, and in a period 0.5-1.5 s after reward. Following rapid onset, these events showed a plateau phase typically lasting up to several seconds. After longer training on an instrumental variant of the task, earlier Plume Events emerged that rapidly followed the learned cue (vibrissa deflection onset).

This initial study discovered spatially organized Plume Events, timed to key behavioral epochs and extending significantly into the neocortical parenchyma. In these data, we employed the contrast created by RhoB to define the vascular wall, an approach that did not allow independent identification of the vessel boundary. Further, our behavioral data from early and later in training were obtained in different cohorts, preventing analysis within-subject and within-vessel. To address these concerns, we next tested a cohort (n = 5 mice), imaging the same PiPA in the three behavioral training sessions, in mice expressing green fluorescence in the endothelial wall (Cdh5-GCaMP8 mice)^95^. We measured and rendered the wall using the gradient profile of the ratio between the red and green signal, excluded the first pixel adjacent to the wall, and tracked this border on a frame-by-frame basis. Data were obtained at higher resolution (pixel length 0.24–1.9 µm; frame rate 3.7–7.7 Hz), allowing more precise granularity in Plume Event detection. To better understand the detailed characteristics of Plume Event emergence, our subsequent analyses initially focused on the PiPA leading into its descent.

This new cohort replicated our initial findings. **Fig. 1b** shows a 3D rendering of a characteristic behavior-timed Plume Event. As shown in this example, Plume Events initiated at one or more specific positions (e.g., at 1 s after cue onset in **Fig. 1d**). This transmission then extended along the vessel surface and, in one or more specific positions, deeper into the neocortex, characteristic of liquid plumes entering surrounding territory. As shown in **Fig. 1b,c,e**, Plume Events were observed in all mice (n = 5) and sessions (n = 15), again showing peaks in amplitude at the peri- and post-reward time points, as in our first two cohorts.

Myriad dynamics are recruited following vibrissa stimulation in SI^96–100^. Most important in this context, another form of vascular dynamics, vasodilation, is well-localized to the time and position of information processing during behavior^101,102^. The many established and hypothesized drivers of functional vascular motion are almost all also leading candidates to create rapid BBB permeability during behavior. Further, stretching the endothelial wall is itself a potential generator of BBB permeability^103,104^. Accordingly, we found that Plume Event probability was increased on trials showing a significant dilation (**Supplementary Fig. 1b**).

To test whether focal dilations are a requisite driver of Plume Events, we analyzed the subset of trials (∼32%) on which no dilation occurred (<0.5% positive or negative change in vessel diameter). **Fig. 1d** shows an example Plume Event from all trials within a session (top), and Plume Events in the same session on only trials without dilations (bottom; Low Dilation trials). While smaller in amplitude, Plume Event emergence without dilations was robust across mice (**Fig. 1e,f**). This finding, that dilations are not required for Plume Events, is consistent with several findings discussed throughout. Interestingly, we also observed that on trials without a detectable dilation, pre-stimulus permeability was higher (teal trace in **Fig. 1e**). Trials without a detectable dilation also occurred predominantly on ‘no response’ trials in which the mouse did not retrieve reward, suggesting a different behavioral state.

The ‘macro’ scale Plume Events identified in our initial study (**Supplementary Fig. 1**) showed spatially specific origin points within the vascular graph depending on time point. We next examined the fine-scale spatial structure surrounding the terminal extent of the PiPA prior to descent into the neocortex. As suggested in the examples (**Supplementary Fig. 1** and **Fig. 1b,d**), Plume Events that showed deeper delivery into the parenchyma were most commonly observed emerging in the axis of the penetrating arteriole. To quantify this effect, we aligned each PiPA to the angle at which the penetrating arteriole ‘dives’ across the neocortical laminae, and created a polar coordinate system centered on the penetrating arteriole (see **Methods**). As shown in **Fig. 1i**, this direction had the greatest probability of Plume Events, both near and far from the vessel, across all mice (**Fig. 1j**).

Our imaging sessions were conducted on the second day of Pavlovian training (P2), the second day of instrumental training (I2) and the eighth day of instrumental training (I8; diagrammed in **Fig. 2a**). When comparing the relative activation maps for each, we found that significantly larger Plume Events were observed on I8, followed by P2 and then I2. These differences are evident in the individual Plume Event maps for each mouse (and in the averages from each; data not shown).

**Figure 2.**
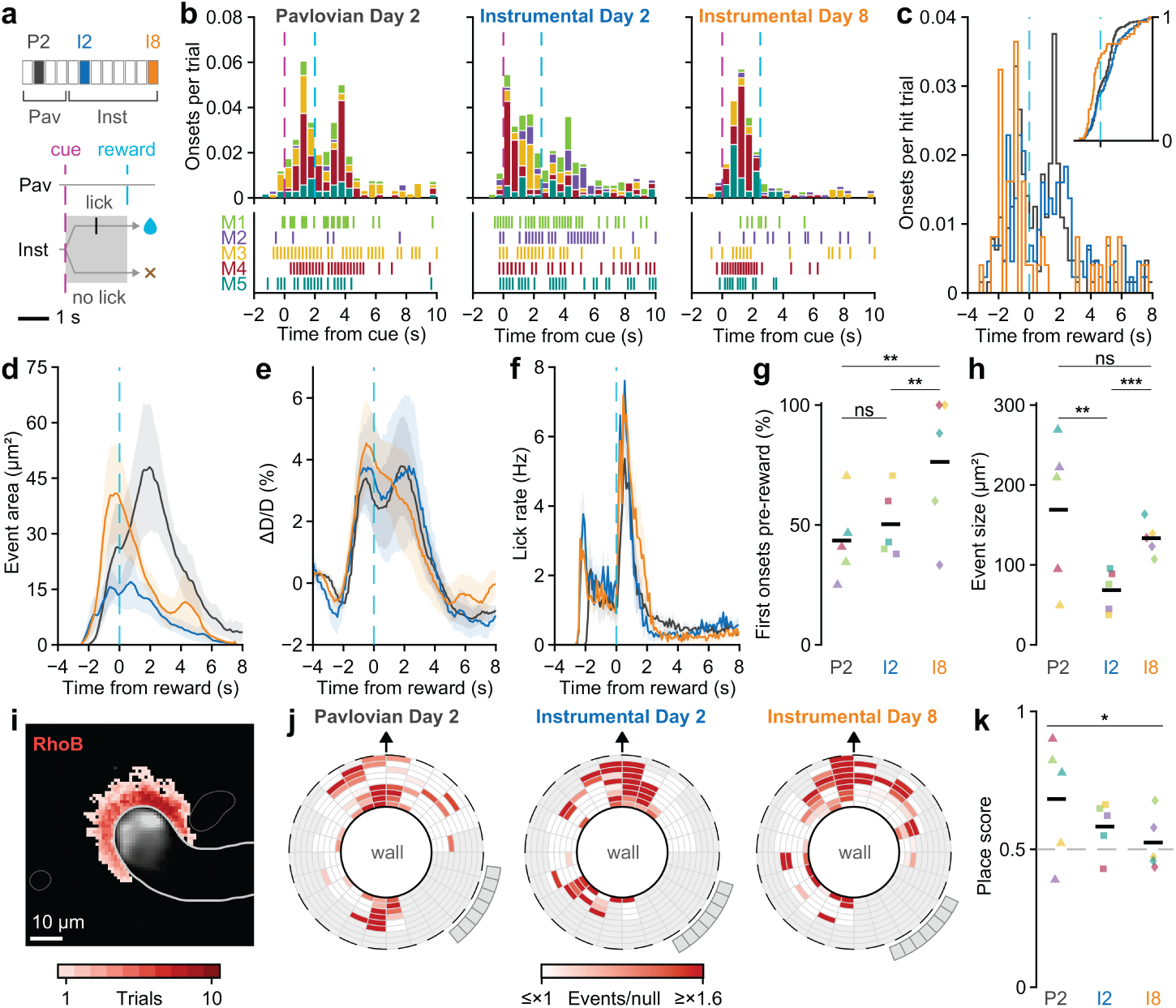
Single-trial Plume Events recur in the peri- and post-reward epochs and are reliably evoked by the cue after sustained instrumental training. **a**, Top: training schedule. Mice were trained on the Pavlovian task for 4 days and then on the instrumental task for 8 days. They were imaged on Pavlovian Day 2 (P2), Instrumental Day 2 (I2) and Instrumental Day 8 (I8). Bottom: trial structure. In the Pavlovian task, water was delivered after every cue. In the instrumental task, water was delivered only if the mouse licked within 2 s of the cue. **b**, Plume onsets per trial relative to the cue on each imaged day, stacked by mouse. Below, onset raster for each mouse. **c**, Plume onsets relative to the reward on hit trials, pooled across mice. Inset shows the cumulative distribution of onsets. **d**, Plume Event area aligned to the reward on hit trials. **e**, Arteriole diameter (ΔD/D) aligned to the reward on hit trials. **f**, Lick rate aligned to the reward on hit trials. **g**, Percentage of first plume onsets on hit trials that occurred before the reward. **h**, Mean Plume Event size. **i**, Number of trials (of 10 example trials) in which a Plume Event covered each pixel in the example session (M1, Instrumental Day 8). **j**, Location of single-trial Plume Events around the vessel wall relative to inter-trial events on each imaged day, rotated with the dive side up. **k**, Place score of single-trial events relative to the average of the other half of trials. A score of 0.5 indicates no relationship to the average location. n = 5 mice. Colors denote individual mice (b, g, h, k) or imaging day (c–f; P2, dark grey; I2, blue; I8, orange). Black bars represent population averages. Shaded regions indicate ± SEM. \**P* < 0.05, \*\**P* < 0.01, \*\*\**P* < 0.001, ns, not significant; mouse bootstrap of a generalized linear mixed-effects model (GLME; g) or a linear mixed-effects model (LME; h), or LME (k).

### Distinct Single Trial Plume Events Occur in the Peri- and Post-Reward Epochs

Plume Events were discovered through trial averaging. The dual-peaked character observed in our trial-averaged data (e.g., **Fig. 1e**) could reflect a single continuous increase, with two peak periods of permeability in the peri- and post-reward periods. Alternatively, as suggested by our initial study (**Supplementary Fig. 1**), these two epochs may represent distinct periods of unique permeability. To address this question, and understand changes in Plume Event size across training, we next analyzed emergence of Plume Events on single trials. All data were analyzed only for successfully detected (‘Hit’) trials, to minimize systematic differences due to behavioral state. Single-trial Plume Events were defined using DBSCAN (see **Methods**).

**Fig. 2b** shows the pattern of onset timings for individual Plume Events on P2. Distinct Plume Events were detected on ∼25% of trials, with distinct onset peaks in both the peri- and post-reward epochs. The pattern of single-trial Plume Events showed a systematic shift in timing across P2, I2 and I8. While I2 showed the characteristic pattern of peaks in the probability of Plume Event onset before and after reward, there was a significant shift to earlier onset times, immediately following the sensory cue, and a decrease in events during the peri-reward period (**Fig. 2b**, *middle*). This shift continued on I8, where single-trial Plume Events were observed to onset almost exclusively prior to reward (**Fig. 2b**, *right*). The relative timing of these events can be observed in the overlapping histogram, aligned to reward time, in **Fig. 2c**.

The trial-averaged data suggested that the shift in Plume Event timing on I2 and I8 reflected an increase in the relative size of these individual events, potentially reducing the observation of later onsets due to continuation of the initial events. To test this prediction, we next calculated the mean area observed cumulatively across the single-trial Plume Events. As shown in **Fig. 2d,g**, I2 and I8 more quickly reach their full areas of transmission following the sensory cue, compared to the later peak in area on P2. On I8, the area occupied is significantly larger before reward than that observed in the P2 or I2 sessions, and on P2 the later, post-reward areas are larger. Notably, the I2 response area was significantly smaller than in either of the other behavioral sessions, a fact observed in all individual mice (**Fig. 2h**). These changes in Plume Events across sessions were not reflected in vasodilation, which remained constant across sessions, or in licking patterns (**Fig. 2e,f**). Further, single-trial Plume Events largely replicated the on-axis localization observed in the mean Plume Events (**Fig. 2j**). In all three sessions, this alignment was significant (**Fig. 2k**).

### Endothelial Cell Calcium Events Mirror the Emergence of Plume Events

Electro-calcium events are a nearly ubiquitous triggering mechanism for the release of behaviorally and biologically important chemical signals. The most notable example is a neural action potential, in which calcium increases in the axonal bouton are essential to neurotransmitter or neuropeptide release^105^. Myriad studies have shown that calcium increases, typically created/measured across minutes to hours, can drive increases in endothelial cell permeability^39,42–44,106,107^. As well, disrupting endothelial cell calcium signaling attenuates the permeabilizing effects of HA on the BBB^84,85^. Further, recent important studies demonstrated in the anesthetized mouse that sensory drive can trigger endothelial cell calcium increases in SI *in vivo*^48,49,52^.

We reasoned that ECCE could drive Plume Events. Our use of the Cdh5-GCaMP8 mouse to systematically track Plume Event distances beyond the vessel wall anticipated this possibility. ECCE were defined using DBSCAN, detecting events lasting > 1 s and encompassing multiple pixels, exceeding a z-score of 3 over their own sliding baseline (see **Methods**). We found that ECCE were robustly driven in our awake task, mirroring the timing of Plume Events. **Fig. 3a** shows an example of a single calcium increase in ECCE at the termination of the axis with the PiPA. This figure highlights the independence of ECCE in this key region relative to task demands, with neighboring ROIs showing active, but independent, dynamics. Much like neural action potentials, we anticipate that ECCE will be discovered to play multiple distinct roles in information processing and vascular dynamics. As one such example, prior studies have shown that ECCE can predict dilations in anesthetized mice^48^. To test whether this relationship held during awake task performance, we tested the prediction that ECCE occurrence predicts subsequent dilation on a given trial. Consistent with prior reports, we found that arteriole ECCE predicted local dilation at a significantly higher rate than those occurring outside this zone or null randomization. However, conjunction of arteriole ECCE and dilation was rare, occurring on only 8% of trials, an effect consistent across all mice (**Fig. 3b,c**).

**Figure 3.**
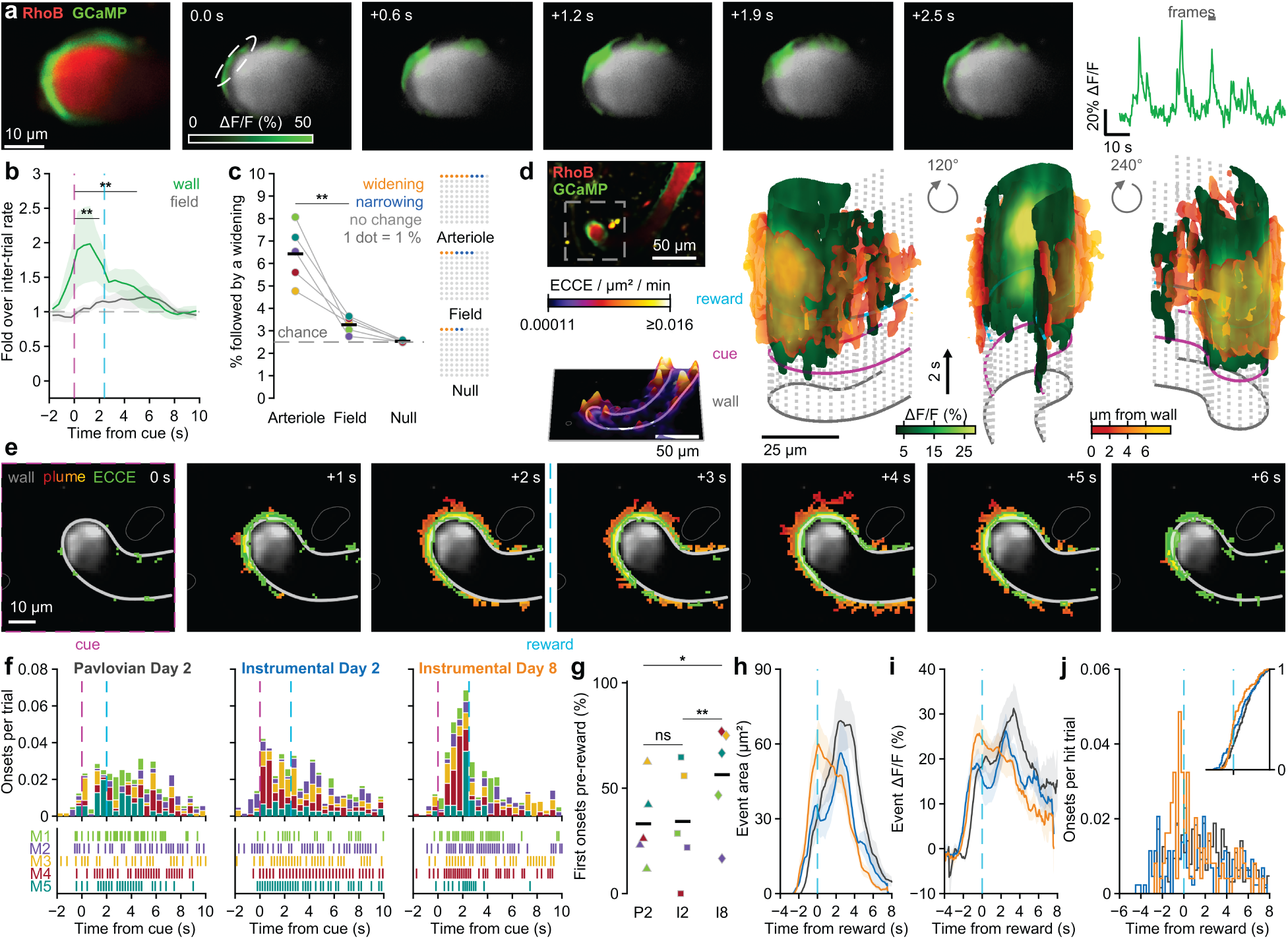
Endothelial Cell Calcium Events (ECCE) at the arteriole wall occur throughout the task and shift to reliable rapid onset with sustained instrumental training. **a**, Example of an ECCE on a penetrating arteriole (M2, Pavlovian Day 4). Left: median image of RhoB-dextran (red) and endothelial GCaMP8 (green). Middle: GCaMP8 ΔF/F at 0.6 s intervals. The dashed outline marks an active region of the wall. Right: ΔF/F within the outlined region; grey bar marks the frames shown. **b**, ECCE onset rate following the cue relative to the inter-trial rate, on the arteriole wall (green) and on arterial vessels in the surrounding field 25–100 µm away (grey). Horizontal bars mark the 0–2 s and 0–5 s windows tested. **c**, Left: percentage of ECCE followed by arteriole widening within 5 s, for ECCE on the arteriole, ECCE in the field, and random times (null). Dashed line indicates chance. Right: the same percentages shown as widening (orange), narrowing (blue) or no change (grey), one dot per percent. **d**, Left: FOV (top) and ECCE density along the arteriole over the session (bottom). Right: three-dimensional reconstruction of the cue-aligned average of trials with an ECCE near the wall (M1, Instrumental Day 8, n = 109 trials). It shows GCaMP8 ΔF/F on the wall (green) and the Plume Event colored by distance from the wall, with time on the vertical axis. **e**, The same trial average as in d at 1 s intervals following the cue, showing the wall (light grey), Plume Event (red to gold) and ECCE (green). **f**, ECCE onsets per trial relative to the cue on each imaged day, stacked by mouse. Below, onset raster for each mouse. **g**, Percentage of first ECCE onsets on hit trials that occurred before the reward. Brackets are marked significant only where the same comparison is also significant for Plume Events (Fig. 2g). **h**, ECCE area aligned to the reward on hit trials. **i**, ECCE ΔF/F aligned to the reward on hit trials. **j**, ECCE onsets relative to the reward on hit trials, pooled across mice. Inset shows the cumulative distribution of onsets. n = 5 mice. Colors denote individual mice (c, f, g) or imaging day (h–j) as in Fig. 2. Black bars represent population averages. Shaded regions indicate ± SEM. \**P* ≤ 0.05, \*\**P* < 0.01, ns, not significant; one-sided mouse bootstrap of a GLME.

To test whether ECCE are positioned to drive Plume Events, we began by analyzing the trial-averaged increases in GCaMP signal. **Fig. 3d,e** show the mean changes in endothelial calcium averaged across all trials for the mouse/session in **Fig. 1b**. This approach reveals a robust increase in calcium in the penetrating arteriole following sensory onset. This increase in mean endothelial calcium fluorescence showed rapid onset (**Fig. 3d,e**) that preceded the mean Plume Event onset. When the mean of each session type was individually compared, the increase in mean endothelial calcium across all trials again preceded Plume Event onset. We next analyzed the distribution of individual ECCE in the three behavioral sessions (**Fig. 3f**). All sessions showed a subset of ECCE with short latencies, occurring ≤ 250 ms after sensory cue onset. Further, paralleling Plume Event onset timing, we again observed a shift to pre-reward ECCE onset on I8, a significant increase in reliability of faster responses compared with P2 and I2.

We next analyzed the specific prediction that individual ECCE predict neighboring Plume Events. **Fig. 4a** shows two examples, the mean time course of ECCE detected on a subset of trials after the sensory cue (top, a P2 session; bottom, an I8 session). As is evident for population data in **Fig. 3**, these ECCE preceded the onset of their corresponding Plume Event. Analyzed across sessions, ECCE showed a uniform distribution in our polar framework. While Plume Events continued to show a bias to the PiPA axis (‘dive’), they also showed a more uniform distribution.

**Figure 4.**
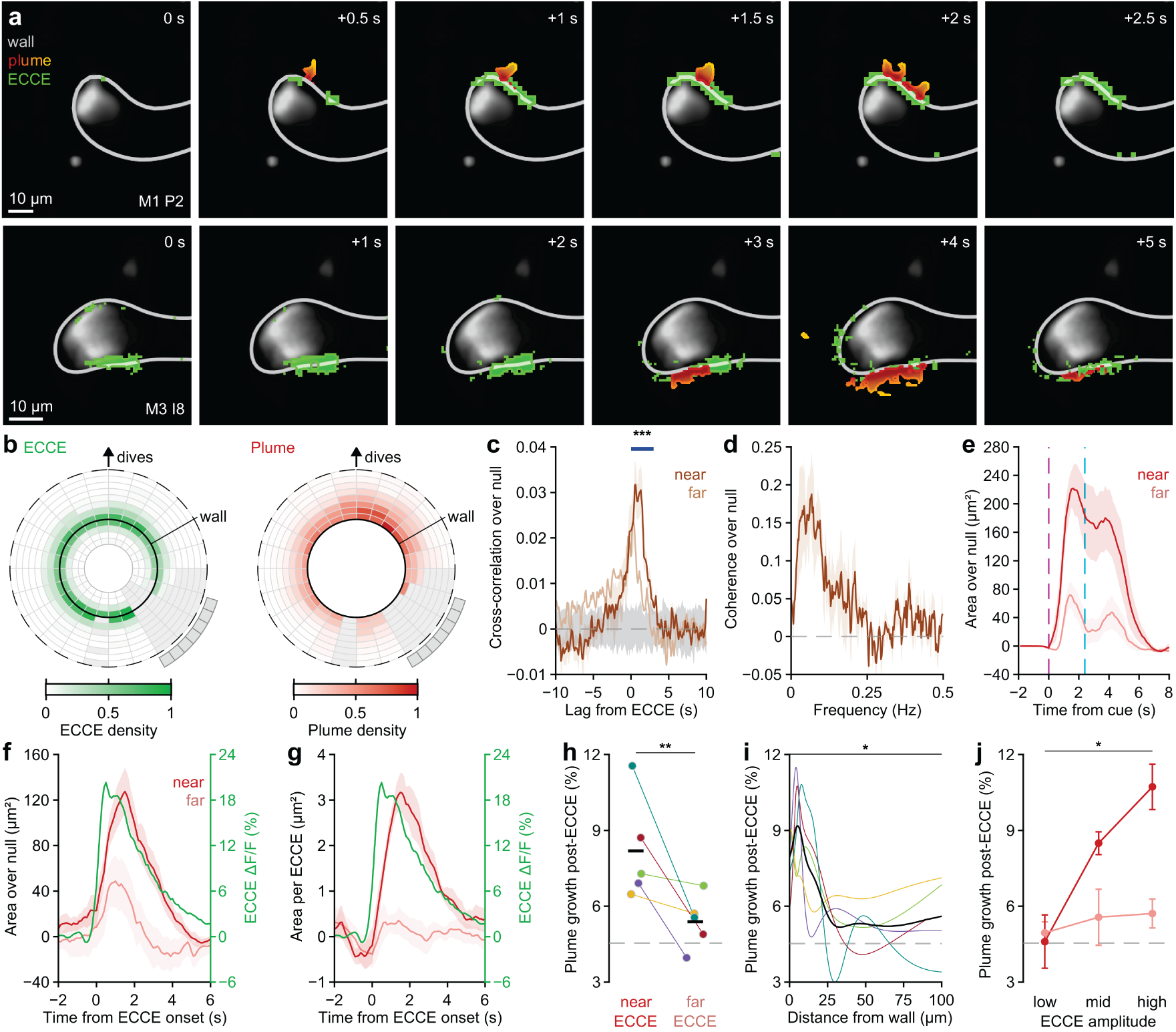
ECCE predict Plume Events that onset at short latency and express <15 µm distance. **a**, Average Plume Event aligned to ECCE onset at one site in each of two sessions. Top: 20 individual ECCE (M1, Pavlovian Day 2). Bottom: 52 ECCE (M3, Instrumental Day 8). ECCE are shown in green and the Plume Event is colored by distance from the wall (light grey). **b**, Location of ECCE (left) and single-trial Plume Events (right) around the vessel wall, pooled across sessions and rotated with the dive side up. **c**, Cross-correlation between the Plume Event signal and ECCE near the wall (within 15 µm; dark) or far from it (more than 25 µm; light), relative to a null made by rotating whole trials. Grey band shows the 95% interval of the null. Horizontal bar marks the window tested. **d**, Coherence between ECCE near the wall and the Plume Event signal relative to null. **e**, Plume area over null following the cue on trials with an ECCE near the wall (red) or only far from it (pink). **f**, Plume area over null aligned to the onset of isolated ECCE near (red) and far from (pink) the wall. Green line shows ECCE ΔF/F (right axis). **g**, Same as in f, measured for each ECCE on its own trial relative to neighboring trials. **h**, Percentage of ECCE followed by Plume Event growth within 5 s, near and far from the wall (n = 2,756 near and 8,194 far ECCE). Lines connect values from the same mouse. Dashed line indicates chance. **i**, Percentage of ECCE followed by Plume Event growth as a function of distance from the wall. Thin lines represent individual mice and the black line the population average. **j**, Percentage of ECCE followed by Plume Event growth by ECCE amplitude tercile, near and far from the wall. n = 5 mice. Colors in h and i denote individual mice. Error bars and shaded regions indicate ± SEM. \**P* < 0.05, \*\**P* < 0.01, \*\*\**P* < 0.001; LME (c), GLME (h), exact one-sided sign-flip test of per-mouse slopes (i), or exact one-sided Wilcoxon signed-rank test (j).

We next tested the prediction that the position of ECCE predicted Plume Events. When ECCE occurred *near* the vessel wall, within 15 µm, a robust cross-correlation was observed, with ECCE anticipating a peak in Plume Event onset probability that was significantly increased for 0–4 s. While a trend was observed, this predictive correlation was not significant when ECCE were > 25 µm distant from the vessel wall. Further, when Plume Events emerged on trials with a near ECCE, they demonstrated a significantly larger area than those on trials with only far ECCE. The spatio-temporal evolution of ECCE-triggered Plume Events can be appreciated in **Fig. 4f–i**, which provide several means of showing the relative size and temporal profile of ECCE and associated Plume Events, near and far. Larger ECCE also predicted greater Plume Event expansion (**Fig. 4j**). In sum, these data indicate that ECCE show similar, but more rapid, temporal dynamics within a task, and similar evolution across trial type in their timing. Further, expression of individual ECCE robustly predicts the emergence of neighboring Plume Events.

### Natural and Directly Stimulated Histaminergic Axon Activity Robustly Predicts Plume Events

HA is a well-established driver of large-scale increases in endothelial calcium signals via endothelial cell HA receptors^108–110^. HA also drives rapid, transient, robust increases in BBB permeability that are dependent on endothelial cell calcium signaling^74–78,82,86^. This rapid increase transforms into suppressed permeability with sustained HA exposure beyond ∼5 min^85,111^. The TMN of the hypothalamus projects HA axons throughout the forebrain, including a dedicated subregion projecting to SI^112,113^. Activity-tagging c-Fos studies indicate that TMN is engaged both when thirst pressure is high and when relieving this state is possible, as in our Pavlovian and instrumental tasks^87^. Further, recent work has indicated a causal role for the TMN in driving learned behavior within day two of a Pavlovian task^114^. We reasoned that these HA inputs may thus drive Plume Events observed in our task, among the many other possible drivers of Plume Events.

**Fig. 5a** shows labeling of TMN somata within the hypothalamus through expression of floxed tdTomato in the histidine decarboxylase (HDC)- Cre mouse. **Fig. 5b** shows the robust expression of TMN axons across the neocortex, including in SI (**Fig. 5c**). Consistent with prior reports73, we further observed that TMN axons frequently pass <2 µm from SI microvessels, including the penetrating arteriole in our two-photon FOV (**Fig. 5d**).

**Figure 5.**
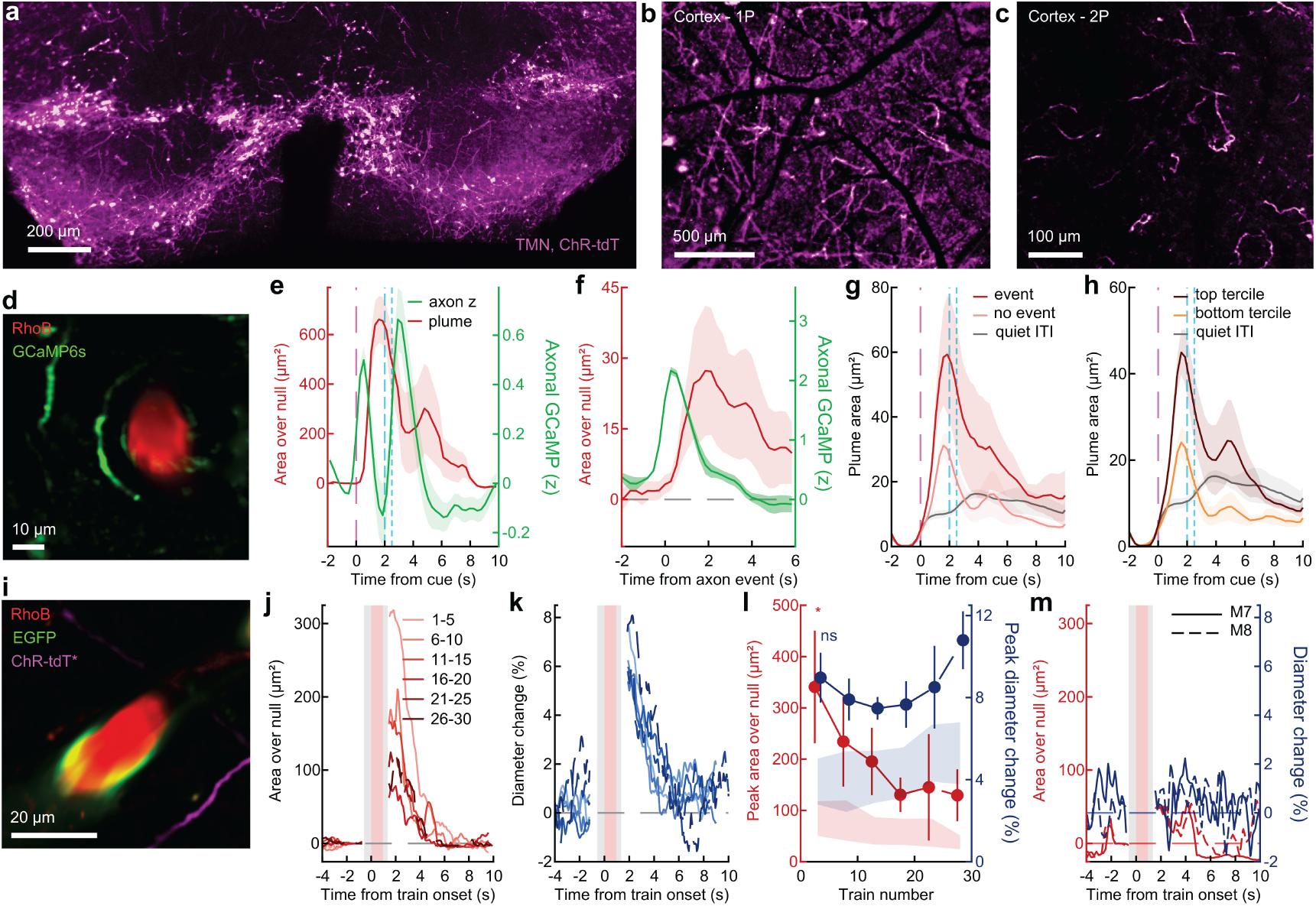
TMN axon activity is rapidly recruited by behavior, immediately leads Plume Events, and is sufficient to drive Plume Events. **a**, ChrimsonR-tdTomato expression in the tuberomammillary nucleus (TMN) of an HDC-Cre mouse. **b**, ChrimsonR-tdTomato in neocortex imaged through the cranial window. **c**, Two-photon image of TMN axons in neocortex. **d**, Example FOV showing a penetrating arteriole filled with RhoB-dextran (red) and a nearby TMN axon expressing GCaMP6s (green) (HDC95, Instrumental Day 8). **e**, Axon activity and plume area over null following the cue. **f**, Plume area over null (red, left axis) and axon activity (green, right axis) aligned to axon events. **g**, Plume area on trials with (red) and without (light red) an axon event, and during quiet inter-trial periods (grey). **h**, Plume area on trials in the top and bottom third of axon activity. **i**, Example FOV from a stimulated mouse showing RhoB-dextran (red), EGFP (green) and ChrimsonR-tdTomato axons (magenta). *Pseudocolored, as tdTomato and RhoB share a channel. **j**, Perivascular dextran area over null aligned to the stimulation train, averaged in sets of 5 trains. Pink shading marks the stimulation and grey shading the frames blanked by the closed imaging shutter. **k**, Arteriole diameter change for the same sets of trains. **l**, Peak dextran area (red, left axis) and peak diameter change (blue, right axis) as a function of train number. **m**, Dextran area (red) and diameter change (blue) averaged over all trains for two mice with no TMN axons in the FOV (M7, solid; M8, dashed). n = 4 mice, 11 sessions, 14 axon ROIs (d–h); n = 6 mice (j–l; n = 4 for trains 21–30). Shaded regions and error bars indicate ± SEM. Dashed magenta and cyan lines mark the cue and reward. \**P* < 0.05, ns, not significant; first 5 trains compared with the remaining trains, two-sided LME.

If TMN HA activity drives the task-related Plume Events we discovered, TMN axons must express a robust, short-latency activation in response to our vibrissa cue. In support of this prediction, TMN receives a robust disynaptic projection from the facial trigeminal nucleus^113^, which mediates the short-latency relay of vibrissal motion^115^. To test this prediction, we trained a new cohort of mice in our task progression and delivered GCaMP6s^116^ driven by floxed expression in HDC-Cre. **Fig. 5e** shows that TMN HA axons (n = 4 mice, 1-2 axon segments in each FOV) are rapidly driven, responding < 250 ms after the sensory cue. We next detected TMN axon calcium spikes that occurred throughout the time series (peak z-score ≥ 3). As shown in **Fig. 5f**, TMN calcium spikes robustly predicted Plume Event onset ∼1 s later. When task trials were analyzed, those with TMN calcium spikes predicted significantly larger Plume Events than those without (**Fig. 5g,h**).

These correlations strongly suggest that TMN HA activation is sufficient to drive Plume Events. To test this prediction, we transduced the cation-passing optogenetic element ChrimsonR^117^ in the TMN. When SI axons were stimulated (n = 6 mice; 1 s, 20 Hz, 5 ms square wave pulse, 625 nm LED) adjacent to our PiPA FOV, we observed robust, transient permeability for the first 5 stimulation trials that decreased with further presentations (**Fig. 5j**). We also observed that TMN axon stimulation drove a robust dilation, a previously reported effect of HA presentation^118,119^. However, these dilations did not degenerate across conditions, remaining robust throughout the session (**Fig. 5k**). The contrast in these two vascular response dynamics is shown in **Fig. 5l**. In 2 mice, no axons were visible within the imaging FOV, and both showed neither increased permeability nor dilation (**Fig. 5m**), confirming that the effects of optogenetic drive (permeability and dilation) were focal to the imaged PiPA.

## Discussion

Here, we describe a novel mode of brain-body coupling, rapid, brief and local BBB permeability timed to relevant behavioral events and TMN axon activity. This behavior-timed permeability is spatio-temporally organized into Plume Events that show predictable differences in size and pattern of emergence. These two classes of Plume Events, peri- and post-reward, were observed in all mice tested, performing either vibrissa-cued Pavlovian conditioning early in training or a well-trained instrumental version of the task. In the well-trained instrumental task, a novel group of Plume Events emerged in response to the trained sensory cue. While we predict Plume Events emerge from many drivers, we tested the role of histaminergic TMN axons, reflecting HA’s impact, these axons’ anatomical apposition to vessels, and TMN’s ‘alerting’ role, one that parallels the occurrence of Plume Events. We found that direct optogenetic activation drove Plume Events, and endogenous calcium spiking in TMN axon segments predicted them.

### Behavioral Tuning

Plume Events discovered here show consistent emergence during specific time windows relative to behavioral epochs. Peri- and post-reward Plume Events were found in all mice tested, across two paradigms with distinct reward contingencies and levels of training. Peri-reward Plume Events were consistently found from 750 ms before to the moment of reward, even when the delay period between sensory cue onset and reward was shifted by 500 ms. Further, sensory cue-related Plume Events were found in 4 of 5 mice tested in the instrumental paradigm. The consistency across mice in these Plume Event onset timings was punctuated by consistent valleys in their probability, most notably the complete absence of any onsets immediately following reward. The consistency of these events, and the emergence of sensory cue Plume Events after more intensive training in the instrumental task context, directly indicate that rapid and local BBB permeability can be spatio-temporally tuned to adaptive behavior.

However, our studies do not shed light on the specific behavioral factors that predict Plume Events. Our initial testing of the State Gate Hypothesis employed two well-established tasks that differed in controlled ways, but these paradigms were not designed to systematically test how peri-reward or post-reward Plume Events may depend on specific behavioral features. Peri-reward and post-reward Plume Events followed discrete external inputs (vibrissa stimulation or predictable water delivery) by ∼750 ms to 1250 ms. A similar peak in Plume Event onset times occurred 750-1000 ms following onset of red light ChrimsonR activation. These data suggest that these Plume Event windows are driven by processes timed to these key task features. These latencies could, however, be related to many other features of task engagement, and could reflect longer (or shorter) latencies to those features.

Specifically, a key factor our tasks did not systematically modulate was across- or within-task predictability, allowing for consistent anticipation. The latency between behavioral trials was jittered pseudo-randomly between 7–12 s (Pavlovian) or 2–7 s (instrumental) after conclusion of the prior trial, meaning mice could anticipate trial onset within a relatively narrow time window. As such, the increase in ‘baseline’ Plume Events in the well-trained instrumental task could readily emerge from such anticipation. Within task, the consistent emergence of peri-reward Plume Events 750 ms to 0 ms before reward presumably reflects such anticipation. As shown across many behavioral study designs, subjects develop a robust prediction of the interval between appetitive sensory cues, including onset of affective response indices prior to stimulation^120^.

### The Central Histamine System as a Dynamic Controller of BBB Permeability across Health and Disease

Here, we show that the central HA system, originating in the TMN, regulates neocortical BBB permeability with fine precision: locally, on rapid timescales, tuned to behavior. As discussed above, TMN activity is recruited during wakefulness, arousal, and motivated behavior, controlling downstream HA tone through extrasynaptic release from widespread projections^53,55,58,68^. TMN axons are anatomically and functionally heterogeneous^112,121^, raising the possibility that the TMN may tune individual regions’ BBB permeability by setting the rate of Plume Events, across behavioral states and sensory inputs. Future work should determine the extent to which TMN-BBB coupling generalizes beyond sensory neocortex, and how temporally and spatially specific it may be. Our results suggest that TMN-BBB coupling may be an important novel mechanism implicated in neurological and/or neurodegenerative diseases such as AD, post-traumatic stress disorder, and addiction. Both TMN dysregulation and chronic BBB breakdown are hallmarks of myriad neurological and neurodegenerative disorders^21–23,72,122^. For example, an overactive TMN during chronic anxiety or addiction could lead to aberrantly high rates and/or poor tuning of Plume Events, facilitating the pathological increases in BBB permeability observed in those conditions^123,124^. Additionally, if Plume Events represent moments of metabolic supply or waste clearance, TMN degeneration could lead to failures in such support, resulting in further declining brain health during AD^125^.

### Circuit Mechanisms

A key prediction of the State Gate Hypothesis is that relay of body information through the Direct Interoception pathway of Plume Events may be triggered by many brain signals. These inputs should, in concept, call distinct locations and potentially create unique types of permeability depending on needs, including informational, metabolic or clearance. Key candidates for such regulation are the many other neuromodulatory systems that mutually target neurons and vasculature^26^.

Plume Events related to behavioral events or TMN axon activity originated on PiPA and, in a subset of experiments where visible, capillaries and venules. These data were obtained at the depth of lower layer I and upper layer II of the neocortical column. Recent studies have shown that PiPA may show distinctly greater permeability than surrounding vascular elements^91^, consistent with the long-standing view that the Virchow- Robin space at the inflection of the penetrating arteriole was a major site for such extravasation. This prior literature was one of our motivations in choosing this focus for our initial testing of the State Gate Hypothesis.

The permissivity of the PiPA BBB may reflect its expression of caveolae^92^, the vesicular membrane compartments typically associated with cross-vascular transmission outside the forebrain. Caveolae are largely absent from capillaries in neocortex^93,126^, likely a major driver of the tight barrier generally observed. The sub-second onset of Plume Events after triggers such as optogenetic drive or TMN spiking (and, potentially, behavioral events) could reflect a rapid release from IV components stored in caveolae: As discussed below, the brief, targeted nature of Plume Events may be best viewed as a paracrine signaling event akin to the release of neuropeptides, consistent with this mechanistic prediction. A potential limitation of this prediction is that we have observed Plume Events originating on capillaries in our FOV.

As discussed below, the molecular mechanisms that may mediate Plume Events are varied and, beyond the observation that TMN axons can drive these effects, currently unknown. That said, Plume Events have highly consistent onset times relative to behavioral events and TMN axon activity, uniformly rapid rise times, and often have a ‘T’ shaped structure of invasion into the parenchyma. All these features suggest that at least a partial mechanism creating Plume Events is opening a non-specific pathway (e.g., a paracellular gate) and not (only) a likely slower molecularly specific transfer mechanism, e.g., transcellular.

## Emerging and Established Views of a Dynamic BBB

The present findings fit well with the emerging view that the BBB, and more generally blood vessels themselves, are dynamically engaged with their environment^31,33^. As Banks has suggested^30^, the BBB may be best understood as an ‘endocrine tissue,’ able to receive dynamic signals and act on its environment, and as Badaut and colleagues^34^ propose, it may be best viewed as an interface, and the ‘Border’ may be a more appropriate metaphor. As recently reviewed, the BBB is increasingly understood to be actively regulated during physiological, as well as pathological, contexts^35^. As discussed in the Introduction, findings showing that, on time scales of tens of minutes, sensory activation can drive the BBB in aligned thalamic and neocortical representations support this general view^36,37^. Further, chemogenetic activation of glutamatergic pyramidal neurons in the hippocampus can also drive enhanced BBB transport within ∼1 h, and the activation and inactivation of multiple endothelial cell genetic programs^29^. The discoveries described here show this dynamism can be truly event-like, operating on the time scale of ongoing behavior, and tuned in multiple ways to the properties of these inputs, and to TMN axon signaling.

### Longer-Time Scale BBB Permeability as the Accumulation of Discrete Events, Not Slow Diffusion

While the stark difference in transmission of intravascular signals to the forebrain initially led to the observation of a ‘barrier,’ myriad studies have shown that in healthy, well-rested brains there can be systematic extravasation. Studies measuring these dynamics, e.g., by imaging the fluorescent dextrans employed here, have almost uniformly been conducted with tens-of-seconds to minutes temporal resolution. The relatively systematic transfer curves, and common views of the BBB, have led to the conception of a net ‘diffusive’ process, created by passive and active molecular level mechanisms. The Plume Events described here suggest instead that these ‘smooth’ curves could also depend on the accumulation of a stochastic process at a faster time scale. This difference has many implications for our modeling of system-wide function.

## Key Next Directions

Several directions for study emerge from the observation of Plume Events. The many questions raised by these new findings posed a challenge in determining the timing for communication of this first description of these results. Next, we comment on limitations of the current findings, and these new directions.

### The Artificial Nature of our Statistical Plume Event Definitions

Plume Events capture the new phenomenon of behavioral and neural event-related permeability by using a statistically defined metric. We developed this analytical approach, and chose the minimally biased DBSCAN algorithm as its anchor, to provide a consistent means of identifying the spatiotemporal structure of our initial net fluorescence findings. However, our methods and this strategy have inherent limitations. Our clustering algorithm approach likely excludes real but smaller events, favoring spatiotemporal persistence of larger, more continuous permeability. Our focus on using an event-related analytical approach in defining Plume Events also has significant limitations, as it places bounds on determining their persistence beyond the trial-related epoch. For this reason, we refrained here from discussing the persistence of Plume Events, instead focusing on their onset patterns and spatio-temporal properties in the initial 1 s post-onset period. This choice in part reflected the relatively low spatial and temporal resolution of our two-photon imaging approach, initially chosen to favor discovery of any new signals as we began testing the State Gate Hypothesis (e.g., higher SNR over a larger window). This imaging approach was ideal for discovery and initial characterization, but was also insufficient for single-trial analyses beyond initial observations. A key next step, requiring faster imaging rates, higher spatial resolution and several forms of enhanced SNR, will be intensive imaging of known Plume Event generation foci to capture their full extent.

### The Shape of Plume Events and Compartments Targeted by Transmission

The Plume Events described here robustly show the rapid relay of signals out of the vessel, timed to behavior and local neural (TMN) activity. However, the compartments targeted by this relay cannot be fully (or even mostly) ascertained from our data for several reasons. Our uniform use of 70 kDa RhoB transmission into the parenchyma had many advantages in this first study of the State Gate Hypothesis, as it provided high signal-to-noise ratio (SNR), an inert, large molecule, and prior studies that have used this relatively well-characterized molecule. The large, inert molecule used here putatively reflects a conservative estimate of transmission during a Plume Event. Smaller, polar molecules such as oxytocin (∼1 kDa, negatively charged) presumably pass more deeply into the parenchyma.

Further, and more generally, using the fluorescence transmission approach, signal in the parenchyma conveys the presence of previously intravascular moieties, not the actual ‘open state’ of the barrier. Response features such as Plume Event onset, spatial position of origin and initial expansion rate are likely all reasonable proxies for key aspects of barrier opening, but other aspects of Plume Events, such as total size and duration, will robustly reflect the ‘stickiness’ of the signal in addition to superfusion of a new indicator. Key next steps will require testing a broad array of possible signals carried in Plume Events, and the use of other signals, such as sensors placed in the parenchyma (e.g., of intravascular signals).

### Other Foci: Different Brain Regions, Capillaries

This first test of the State Gate Hypothesis was conducted in primary somatosensory vibrissa barrel neocortex. This choice reflects extensive existing knowledge of the neural and vascular dynamics of this system^96,97,99,100^. The dynamics discovered in SI, and the principles and circuit mechanisms elucidated there, have shown transferability to humans and to other neocortical areas^127^. As such, this choice made initial sense, and there is optimism that the new discoveries made here are not idiosyncratic to the area. However, this evolutionarily ancient body map could have evolved mechanisms for privileged access to body state. This kind of interoceptive alignment has recently been studied intensively in insular regions, primarily through alignment of action potential relay systems (referred to here as Indirect Interoception)^128–130^. However, hormones such as glucagon-like peptide-1 and insulin also act within the insula^131,132^. Whether such relay occurs with the rapid, transactional dynamics observed in SI Plume Events remains to be seen. More generally, how the findings here apply to other forebrain structures—e.g., to aligned striatal representations—requires further study.

For this first study testing the existence of Plume Events, we also chose a systematic focus, PiPA. This choice reflected the well-studied behavior of these vessels during functional hyperemia (particularly in SI) and a consistent anchor for cross-experiment comparison. Further, these vessels show substantial expression of caveolae, vesicles that can mediate transcytosis of several molecules, including insulin and leptin^133,134^, and mice lacking caveolae have impaired transcytosis of aminopeptidase P^133,135^. While used throughout the body for such transmission, caveolae are not present in the rest of the neocortical vasculature. Their absence has been proposed as a major distinction of the forebrain BBB. The behavior- and TMN-driven Plume Events found here could act through triggering release of these vesicles. A rapid, synaptic-like release would satisfy the speed and temporal precision of Plume Events, as they could accumulate larger intravascular molecules at a relatively slower rate, having them prepared for discrete, rapid release events. If so, this transactional and discrete release process could also provide the consistency in rise time, reflecting a biophysically consistent pro-fusion time period for caveolar cargo release. Alternatively, paracellular transmission could also satisfy the temporal precision and speed criteria, but the size of molecule we observed to show relay in Plume Events (70 kDa) is inconsistent with computational modeling of such transmission^136,137^. That said, unique structures such as tricellular junctions could provide unique gating of larger molecular structures^138,139^. If synaptic-like caveolae release is the exclusive generator of Plume Events, then other vessel types should not display this behavior. While our sample was smaller, we did observe Plume Events originating proximal to other vessel types.

The novel discrete form of brain-body coupling described here has many potentially important functions, including serving as Direct Interoception, unmediated access to the full spectrum of body signals by local forebrain circuits. Such Direct Interoception could inform state-tuned learning and choice. Plume Events could also have myriad roles in brain health, including spatiotemporally timed delivery of new metabolic supply and focal waste clearance, when and where waste is generated. We next review the motivation for these studies and these implications.

## Why Plume Events Matter

We tested the ideas presented here for several reasons. A primary motivation is the hypothesis that vascular dynamics of several forms could play an active role in computations underlying adaptive behavior (the ‘Hemo-Neural’ Hypothesis)^26^. This proposal reflects many intuitions about computation underlying behavior, including the fact that the vasculature is a dense and active network intermingled with established forebrain circuits. This network can show myriad forms of dynamics, including, in addition to the fast BBB effects described here, fast mechanical signaling (e.g., dilations and constrictions), changes in flow rate (which can impact factors such as temperature), and fast calcium signaling (e.g., both events and waves)^48,49^. Fast mechanical signaling, local vasodilations and constrictions, occurs in brain areas and specific positions (e.g., neocortical columns) where information processing is crucial to behavior, suggesting it could play a role in modulating local circuits, e.g., in the allocation of attention. Perhaps the strongest argument for an active vascular role in computation is its juxtaposition between neural networks and high-dimensional body signals, the two kinds of signals most needed for adaptive behavior. Integration by endothelial cells and more generally the BBB could allow vessels to make joint computations across these systems, for example determining the risk of permeability and the need for information delivery to the forebrain, and deploy dynamics accordingly.

## A New View of Direct Interoception

The high-dimensional signaling flow in the vasculature is precisely the information needed, in all its detail, for the crucial choices and learning that forebrain neural systems evolved to address. The forebrain provides the ability to learn a complex representation of potential best actions, an ability limited by the quality of information about the problem space of organismal needs it is solving, one that exists at many time scales (e.g., immediate, daily, global). While interoceptive information can reach the forebrain through the pattern of action potential firing in real-time by many ‘classical’ pathways, such pre-formatted signals are, as with all representation, an abstraction of specific features. In contrast, the molecular reservoir in the vasculature includes, at any given moment, updates from almost all body systems, from the parallel intelligence of the immune system to ongoing signaling in myriad organs. This information stream is updated on a seconds-to-minutes time scale. This molecular signaling stream is also, in concept, equally available to access from any point in the forebrain, as all are near the dense vascular system.

The BBB is canonically seen as providing diffusive information to the forebrain to inform action. This slower signaling view has evolved from the general observation of the unique limitations to spread in the forebrain, and has been supported by studies taking data regarding transmission at terminal end-points, or at tens of seconds or minutes time scales. As discussed above, a long history of studies has shown that functionally relevant and localized processing can influence this permeability, as can shifts in body state such as circadian rhythms^140–142^. Here, we show the BBB also possesses a stochastic, rapid and transactional mode of local, fast communication, Plume Events. These instances of potential communication are consistent with a constant— if highly focal—communication modality, more akin to action potential signaling. Local networks may call for this information when, as mentioned above, the risk of exposure to the forebrain is worth the reward in the information gained. Specifically, Plume Events may be best viewed as a kind of ‘paracrine’ signaling, a discrete moment of communication that could have a spatiotemporally diffuse impact for up to seconds (or longer) after release^32^. Plume Events differ from a classic paracrine formulation by conveying a wide spectrum of information signals (e.g., as opposed to a single neuropeptide release). Perhaps Meta-Paracrine signaling.

## Role for Rich Information in Plume Events: Learning

Arguably, the greatest feature of forebrain function is its capacity for learning complex functions that combine internal and external variables with ongoing reward or punishment. Rapid local BBB signals at crucial learning moments—when a crisp feedback signal has occurred indicating the relevance of specific ensembles recently active at a location in the brain—could provide such signaling. As emphasized in Swissa et al.^38^, the sensory-driven permeability to albumin they observed, occurring on a minutes time scale, could play this role. The ‘snapshot’ of information from body signals could, in concept, teach those local ensembles that possess the signaling mechanisms that were just successful in driving a reward (correlated with it) to be more sensitive to the next upcoming state, better able to recognize it even without a signaling event through the typically weak signal relay from vessel to brain.

## Role for Rich Information in Plume Events: Real-Time High-Dimensional Choice

The forebrain applies the complex patterns it learns to making optimal real-time decisions. Such choices require information about the needs of the system at many time scales, immediate, daily and global. The brief snapshot of body state provided to a circuit engaged in a task could, during decision making, provide information that enhances the specificity and criticality of judgments made. Weighing the decision between staying to protect young or forage for a drink can be viewed as a simple binary choice, but can be deeply informed by a wide range of variables. The recent history of predator sounds and smells, weather conditions and time of day that might favor predator behavior learned across a lifetime of observation, the rate and volume of pup calls that could increase predator risk, the sufficiency of other adult conspecifics to help protect young, and nearly endless other external sensory variables could contribute to this decision making. A similarly nearly endless set of internal variable signals should also determine choice: Is risk of dehydration acute, could drinking improve the quality of brain function and judgment in the next several hours, a riskier time of day (e.g., getting a drink midday versus at crepuscular sunset), are oxytocin levels high suggesting that calculations of pup need are high in their own right? Is the act of rumination itself a cost in the value of a decision? At their best such decisions are not simple verbally framed binary choices: In humans and in all non-verbal creatures, many such decisions will be best made by fitting a complex optimization function to drive the abstract sequences needed to implement such a choice (the route to the river, running and pausing in irregular non-predictive intervals). Put another way, optimal intuitive choices whose variable space can go far beyond those easily cognitively stated require complex sensory and interoceptive variable matching to many time scales of signals.

Plume Events are a potentially ideal source of such rich information at times when large-scale variable integration is crucial. Release of endocrine factors that can select ensembles—those that past experience indicates are best for the task, and those that perhaps have different settings for optimal recruitment through multiple drivers (mild oxytocin with low vasopressin sensitivity but a strong non-linearity to vasopressin)—is crucial. Factors in Plume Events targeting known controllers of ensemble selection, such as local axon segments from long-range modulatory systems known to create attentional specific state allocation (local HA, acetylcholine or norepinephrine inputs) or interneurons should be key targets.

## Role for General Effects of Plume Events

While the specificity of Plume Event modulation through the rich signal depth is a strong argument for this local form of brain-body coupling, vasculature contains myriad factors that could more generally enhance function in local circuits, if introduced in a limited and targeted way. As one such example, Friedman and colleagues^38^ recently showed that focal albumin delivery can enhance synaptic plasticity, and proposed that a corollary amplification of learning could reflect the activity-dependent BBB permeability that has been repeatedly shown to occur on tens-of-minutes time scales.

## Health Benefits: New View of Local Metabolic Supply

While local vasodilation (e.g., of PiPA components) maps well to local brain regions engaged by task demands or attentional allocation, many kinds of evidence show that this response is not well correlated to metabolic need^27^. This finding is often ignored (e.g., in commonly used textbooks) as it is unintuitive. In other body systems (e.g., skeletal muscles), changes in vascular supply are well-matched to metabolic need. Further, the high consumption rate of the brain suggests the need for optimization, even in the context of high levels of ambient metabolic supply due to dense vascularization.

Plume Events may provide an alternative pathway for tailored, highly focal metabolic supply. Transient BBB permeability could deliver glucose and other micro- and macronutrients necessary for neural function to the circuits engaged by specific existing task demands. As shown in our TMN findings, functional hyperemia is not inherently required to generate Plume Events, though there may be many important interactions between these two forms of vascular dynamics. As such, focal metabolic supply could be delivered by Plume Events to active circuits in ways that are better correlated to need, providing an answer to the puzzling disconnect that has led so many textbooks to restate a disproven view about functional hyperemia.

## New View of Local Clearance

By a similar token, Plume Events could also create a local clearance mechanism, localized to the time and place of dynamic waste creation. In addition to dynamic mechanisms such as focal dilation ‘pumping’ of glymphatic clearance^143–146^, Plume Event-based clearance could provide tailored extraction. This mechanism should be most effective for generators adjacent to vessels. In general support for this proposal, the BBB is known to serve as a major sink for many kinds of waste clearance^20,28^. As one example, ∼25% of Aβ clearance is believed to occur through the BBB^147^.

### New View of BBB Mechanisms in Brain Disease

The State Gate Hypothesis proposes that Plume Events are a powerful, tuned means of linking arousal and actions to body state to drive optimal healthy behavior. Alterations in HA, and HA pathway signaling, and the BBB, in neurological and psychiatric disease could either entirely nullify Plume Events or flatten their sensitivity. Under this outcome, individuals would lose the ability to align normal body needs to forebrain computations around choice. Addiction is, at its core, an advanced system for such loss. Similarly, changes in BBB permeability could contribute to anxiety. Failure to connect to body states through loss of, or change in, Plume Events could facilitate ruminative behavior, focused on external drivers such as social interaction. Conversely, catastrophizing illness state is often regarded as emerging from too present an awareness or central representation of body state. Whether Plume Events might drive conscious awareness of body state, or explicit memories that replay prior realizations of this state, is an intriguing if entirely open question.

### New Approaches for Drug Delivery

Transmission of molecular therapeutics across the BBB is the focus of intensive study and invention, with a focus on transient disruption of vessel integrity or the invention of new ferrying molecules^148,149^. The mechanisms driving Plume Events are not known, and the molecules they can transmit have not been determined. The fact that a large, non-biological, neutrally charged molecule (70 kDa RhoB) can be relayed through Plume Events suggests that a broad spectrum of agents may also be transmitted through this mechanism. If useful pharmacological agents can be relayed this way, behavioral and/or histaminergic modulation during IV drug presentation could be a powerful, novel approach to delivery.

### Plume Events as a Pathway for Brain-to-Body Relay

The brief periods of transmission from the vessel to the brain found here may also reflect moments of brain-to-body relay. Such transmission could provide a unique chemical signaling pathway from the forebrain, relaying both specific (e.g., endocrine) signals and set-point amounts of brain-derived molecules, thereby reflecting overall activity levels. By a Plume Event mechanism, such transmission would have spatio-temporal tuning, providing the body with a snapshot not just of overall brain function, but specifically of the signals carried in parts of the forebrain engaged in information processing.

## Methods

### Animals

This study was performed in accordance with the recommendations of the Guide for the Care and Use of Laboratory Animals of the National Institutes of Health. All procedures were approved by the Institutional Animal Care and Use Committee of Brown University (Protocol 26-10-0003). Mice were between 3 and 9 months of age and were individually housed after surgery. Food and water were provided *ad libitum* outside of behavioral training, and animals were housed on a 12 h light/dark cycle with all experiments occurring during the dark (active) cycle. Plume Events and endothelial calcium were imaged in five Cdh5-GCaMP8^150^ transgenic mice (#033342, The Jackson Laboratory, Bar Harbor, ME; 4M/1F), which express GCaMP8^151^ in the vascular endothelium^95^. In a separate cohort, TMN axon activity was imaged in four HDC-Cre^152,153^ mice (#021198, The Jackson Laboratory; 3M/1F) expressing an axon-targeted, Cre-dependent GCaMP6s^116,154^ in the tuberomammillary nucleus (TMN). Optogenetic stimulation was performed in eight HDC-Cre mice (6M/2F) expressing the opsin ChrimsonR^117^ in the TMN. Experimenters were not blinded to experimental conditions during data collection or analysis.

### Surgical procedures

Mice were anesthetized with isoflurane (3% induction, 1.5% maintenance) vaporized in oxygen at 1.5 L/min, following procedures described previously^155,156^. Before surgery, mice received intraperitoneal dexamethasone (0.2–0.6 mg/kg) and subcutaneous extended-release buprenorphine (0.5– 1.0 mg/kg). Body temperature was maintained with a heating pad, and an ophthalmic ointment (white petrolatum and mineral oil) was applied to the eyes. The incision site was numbed with subcutaneous 1% lidocaine (3–5 mg/kg) and sterilized with betadine and 70% ethanol, and the skin atop the skull was retracted. The fascia was dried and removed with a 0.5 mm carbon steel drill bit (19007-05, Fine Science Tools, Foster City, CA) and the outer skin margins were secured with tissue adhesive (Vetbond, 3M, St. Paul, MN). A custom-machined titanium head-bar was then adhered to the skull with dental cement (C&B Metabond, Parkell, Edgewood, NY). A 3 mm craniotomy was centered on the vibrissa representation of the primary somatosensory barrel field, at AP −1.25 mm, ML 3.5 mm from bregma^157^. The bone was thinned and removed with a 0.3 mm tungsten carbide drill bit (H71.HP.003, Komet Dental, Lemgo, Germany) following established methods^158^, and the window was sealed with a 3 mm, #1 thickness round coverslip (CS-3R, Warner Instruments, Hamden, CT). For the TMN cohorts, adeno-associated virus (AAV) was injected bilaterally through burr holes at AP −2.7 mm, ML ±0.9 mm, DV −5.2 mm from bregma^159^. Virus was infused at 50 nL/min, 300 nL per site: AAV1-Syn-FLEX-rc[ChrimsonR-tdTomato] (Addgene #62723-AAV1) for optogenetic experiments and AAV5-hSynapsin1-FLEx-axon-GCaMP6s (Addgene #112010-AAV5) for axon imaging at the titer supplied by Addgene (Watertown, MA). The cranial window was implanted in a second surgery approximately four weeks later. In the optogenetic cohort, the endothelium was labeled by retro-orbital injection of the X1.1 capsid^160^ carrying CAG-EGFP (Armamentarium Vector Core, California State Polytechnic University, Pomona, CA), 1 × 10¹¹ vg in 100 µL of sterile saline. Mice were given at least one week to recover from surgical procedures before habituation. They were then gradually acclimated to head fixation of increasing duration (15, 30 and 60 min) on the days preceding the onset of experiments.

### Behavioral training

Three days before the onset of behavioral training, mice were water restricted to ∼90% of their baseline body weight and never allowed to fall below 85%^161^. Water was not available in the home cage, other than a top-off given 1 h after the end of each session. Mice were trained on a vibrissa-cued reward task for four days of Pavlovian conditioning followed by eight days of instrumental conditioning. The cue was a 20 Hz train of ten vibrissa deflections (500 ms), each a 25 ms asymmetric cosine with a fast-rising phase and a slower return, delivered by a piezoelectric bender (NAC2228, CTS Corporation, Lisle, IL)^162,163^. In the Pavlovian task, water was delivered 2 s after every cue; in the instrumental task, water was delivered at ∼2.5 s only if the mouse licked within 2 s of the cue (the commit window), and otherwise a dummy solenoid produced an identical click with no water. Each reward was 4 µL of water, and each session ran for 40 min, giving approximately 150 trials. The cue was withheld until 2 s had passed with no lick bout, so that no cue in any analyzed session fell within 2 s of the onset of a lick bout. Otherwise, the inter-trial interval was drawn from a uniform distribution (7–12 s in Pavlovian, 2–7 s in instrumental), giving cue-to-cue intervals of 13.5– 18.5 s and 11.75–16.75 s, respectively. Imaging of Plume Events was performed on Pavlovian day 2 and instrumental days 2 and 8. Behavior was acquired with Bonsai^164^ through a Teensy 4.1 (PJRC, Sherwood, OR), which logged the cue piezo, the reward solenoid, the failure solenoid, a capacitive lick sensor, and a behavioral camera. A second Teensy 4.1 generated the piezo waveform at 5 kHz through a 12-bit digital-to- analog converter, triggered by Bonsai at cue onset. Behavioral and physiological monitoring followed approaches described previously^165,166^. Signals were placed on a 1 kHz grid shared with the imaging frame counter, binarized and downsampled to 10 Hz. Solenoid and piezo events were debounced with a 1 s dead time, and failure pulses within ±1 s of a reward were discarded. Licks were scored as sensor occupancy in 100 ms bins and a lick bout as a rising edge. One trial was defined per cue, and the last cue of each recording was dropped. In Pavlovian sessions a hit is a trial on which the mouse drank; in instrumental sessions a hit is a trial on which it licked within the commit window and drank. Trials on which the previous trial’s water was left undrunk and the mouse licked in the current trial were marked contaminated and excluded from the hit/miss classification.

## Two-photon imaging

Before each imaging session, mice received 50–100 µL of 0.75% (weight/volume) 70 kDa rhodamine B-dextran (D1841, Thermo Fisher Scientific, Waltham, MA) dissolved in sterile saline. The Cdh5-GCaMP8 cohort was injected by tail vein without anesthesia; the TMN axon and optogenetic cohorts were injected retro-orbitally under brief isoflurane anesthesia. Mice were then head-fixed on a locked wheel. The retro-orbitally injected cohorts were given 30 min to wake up before data collection^167,168^, while the Cdh5-GCaMP8 cohort was imaged immediately after injection. Imaging was performed in the awake animal^169^ on a two-photon microscope^170^ (Bruker, Billerica, MA) running Prairie View, with a CFI75 LWD 16X 0.8 NA water-immersion objective (Nikon, Melville, NY). Excitation came from a Chameleon Ultra II Ti:sapphire laser (Coherent, Santa Clara, CA) tuned to 920 nm, at 15–25 mW measured at the sample. Red and green emission was separated with the standard Bruker filter set: a T565lpxr long-pass dichroic with ET525/70m-2p and ET595/50m-2p bandpass filters (Chroma). The red channel collected rhodamine B-dextran and the green channel collected Cdh5-GCaMP8. In every field of view, imaging was centered on a pial arteriole from its branch point to the point at which it descended into the neocortex (the pial-to-penetrating arteriole, PiPA) at the layer I–II border, 50–100 µm below the pia. The effective frame rate was 3.73–7.72 Hz and the median pixel size per cohort was 0.13–0.71 µm.

## Optogenetic stimulation

TMN axons were stimulated through the cranial window with a 625 nm LED (M625F2, Thorlabs, Newton, NJ) coupled to a 400 µm core, 0.39 NA optical fiber (FT400UMT, Thorlabs) and driven by a Cyclops LED driver (Open Ephys). The fiber was positioned essentially flush with the glass, at about 45° to the window and beneath the objective. The illuminated spot was therefore approximately the diameter of the fiber core, elongated along one axis by the oblique angle. Each session delivered ∼30 stimulation trains, each 20 pulses of 5 ms at 20 Hz (0.955 s)^171,172^, separated by ∼57–59 s over a 30 min recording. Optical power at the fiber tip was 4–5 mW, approximately 30 mW/mm² across the fiber core. The PMT shutters were closed 100 ms before the first pulse of each train and reopened 100 ms after the last, blanking 1.2 s around each train. In two of the eight mice, no TMN axons were present in the field of view.

## Histology

Following the conclusion of imaging experiments, a subset of animals was deeply anesthetized with an intraperitoneal overdose of Beuthanasia (≥150 mg/kg) and transcardially perfused with PBS followed by 3.7% formaldehyde, diluted from a 37% stock (F8775, Sigma-Aldrich, St. Louis, MO). Brains were extracted, drop-fixed for 24 h and transferred to PBS for long-term storage. Fixed brains were cut into 50 µm coronal sections on a VT1000 S vibratome (Leica Biosystems, Deer Park, IL) and held in 24-well plates until they were mounted on microscope slides (12550123, Fisherbrand, Thermo Fisher Scientific, Waltham, MA) and coverslipped with DAPI Fluoromount-G (0100-20, SouthernBiotech, Birmingham, AL) under #1.5 coverslips (48393-241, VWR, Radnor, PA). Sections were imaged on an Eclipse TE2000-E inverted fluorescence microscope (Nikon Instruments, Melville, NY) fitted with an X-Cite Xylis II broad-spectrum LED illuminator (Excelitas Technologies, Pittsburgh, PA) and a Photometrics X1 cooled CCD camera (Teledyne Photometrics, Tucson, AZ), through Plan Apo 4× (0.20 NA) and 10× (0.45 NA) objectives and processed in Fiji^173^.

## Data Analysis

### Image registration

Image stacks were corrected for x–y motion with a rigid, translation-only registration based on discrete Fourier transform phase correlation^174^. The reference was the mean of the red and green projections over the last 50% of frames. Shifts were estimated with 100× upsampling and applied as whole-pixel translations; a frame whose shift departed by more than 3 px from the 11-frame moving median was moved by that median instead. The first frame was dropped and the stack cropped to the valid region.

### Arteriole diameter

Per frame, the cross-sectional area A of the penetrating arteriole was measured by Radon segmentation and converted to an area-equivalent diameter, d = 2√(A/π). The Radon transform of a box around the vessel was taken at 0–179°, the sinogram thresholded at half-maximum and inverse-transformed, and the resulting image thresholded at the Otsu baseline plus 0.2 × (99.5th-percentile peak − baseline). The area of the largest region was taken as A^175,176^. A rolling mean of round(1.3 × frame rate) frames was applied before the Radon step, and despiking and a 1 Hz low-pass filter afterwards. Low Dilation trials were selected by dropping, one at a time, the trial that pulled the set’s mean diameter (as a percentage of each trial’s own pre-cue mean) furthest from zero, until that mean stayed within ±0.5% at every lag from 0 to 5 s. At least 10 trials were retained. Ten of 15 sessions yielded such a set (median 52 trials, range 25– 135).

### Wall tracking

The arteriole wall was drawn once by hand on each session’s cue-aligned trial-averaged movie and then moved at every lag by a single signed offset applied uniformly along the outward normals of that seed. That offset preserved the seed’s shape and was measured from the seed each time rather than accumulated across lags. The offset was driven by the outer edge of the dye lumen at a fixed pre-cue half-maximum level, with a bounded contribution from the green endothelial profile (a cross-correlation shift within ±2 px, weighted at most half as strongly as the red). The resting wall was the union of the tracked positions over the pre-cue lags. Every subset average re-tracked the wall on its own movie. For single-trial analyses the resting wall was instead pushed outward frame by frame by half of that trial’s increase in penetrating-arteriole diameter over its own pre-cue mean (the carved wall). Area was counted only beyond max(2 µm, 1.5 px) of it.

### Plume Event detection

Plume Events were detected on the red channel after smoothing with a 0.7 px Gaussian and z-scoring each pixel against its own mean and standard deviation over the whole recording. For trial-averaged analyses, trials were aligned to the cue, averaged from −1.9 to +10.0 s on native frame times and smoothed with a centered moving mean of round(0.7 × frame rate) frames. The search band comprised pixels up to 15 µm outside the tracked wall at each lag, inside a hand-drawn region of interest and outside exclusion zones drawn over bright off-vessel structures. Supra-threshold pixels were grouped in (x, y, t) by DBSCAN^94^ (eps of 3 px in x and y and 1.25 s in t, minPts 5), and a Plume Event was defined as a cluster that touched the inner 2.5 µm of the band at some lag (the wall-origin gate). For the cohort statistics, the threshold at each pixel and lag was the upper limit of a 95% confidence interval built from the random-alignment null (below), pixels above threshold at every lag were not searched, and the Plume Event was the largest admitted cluster, trimmed at each lag to its largest connected component. Excess area at a lag was the real area minus the mean area of the null draws at the same lag. For the montages, each pixel’s rise over its own pre-cue mean was thresholded at the 99th percentile of the in-band baseline rises of the same average, and every wall-touching cluster was kept. For the spatial analyses, the band and the wall-origin gate were opened to the whole region of interest. Density was the supra-threshold area divided by the searchable area, in 2 µm bins of distance from the wall or in perivascular (≤ 15 µm) and parenchymal (> 15 µm) zones, averaged over the reward window [rc − 0.75, rc + 2.25] s, where rc is the session’s median cue-to-reward delay; for the dive-side map, each vessel was rotated so that the side on which it leaves the imaging plane points up and mirrored so that its parent segment lies to the right. On single trials, each pixel’s mean over the pre-cue frames (−1.97 to −0.26 s) was subtracted, and the threshold was the 99th percentile of those pre-cue increases pooled over the session’s trials within the band, which started at the carved wall. Supra-threshold voxels were grouped by DBSCAN with eps of 1.5 px and 1 frame and minPts 3, and a cluster was kept if it contained at least 5 voxels and had a voxel within 2.5 µm of the carved wall in its first 0.5 s. A single-trial Plume Event was a kept cluster at or above the 95th percentile of the largest clusters found in that session’s quiet inter-trial windows (below).

### Null models

Chance was estimated by resampling within each session, so that every null carried that session’s own dye kinetics, motion and vessel geometry. The primary null was a random-alignment null: 100 trial averages per session, each built from the same number of trials as the real average but with every trial anchored at a time drawn at random from anywhere in the recording, and each passed through the identical threshold, clustering and gates. Because the anchors could fall anywhere, the draws include real cue and reward periods. The cohort detection threshold was built from such averages, as their mean plus t(0.975, lags − 1) standard errors at each pixel and lag. One-sided P was computed as (1 + number of draws ≥ real) / (1 + number of draws), whose smallest attainable value with 100 draws is 0.0099, and drawn null bands are the central 95% of the null of the same estimate. Two restricted versions addressed specific alternatives. A quiet-anchor null confined anchors to inter-trial time at least 7 s after the previous reward, with no cue, reward or failure in the following 1 s. A lick-free null confined anchors to times with no lick-bout onset in the previous 2 s and no lick in the previous 1 s, matching the cue-withhold rule, which the real cues satisfy on 93.1–100% of trials per session. For single-trial events, chance came from 200 quiet windows per session, drawn without replacement at least 10.8–11.6 s after the previous cue and at least 9.0 s after the previous reward or failure, with no cue, reward or failure in the following 1 s, and passed through the identical detector at the same pixel threshold. For the place score, spatial chance was obtained by sliding each event’s footprint along the resting wall in 1 px steps all the way around, preserving its arc length and distance from the wall; a slide was valid if at least 90% of the footprint landed on support, and at least 10 valid slides were required. The score is the percentile of the real position’s mean reference value among the slides, where the reference is the occupancy of the trial-averaged Plume Event computed from the other half of trials, split by odd and even cue number. For model-based tests, uncertainty across animals was assessed with a cluster bootstrap in which mice were resampled with replacement and the model refit (2,000 resamples for a linear model, 400 for a generalized linear model, seed 42); with five mice this gives 126 distinct resamples.

### Endothelial calcium events

ECCE were detected on the green channel over the full frame. Each frame was pre-smoothed with a two-dimensional Gaussian of ∼0.7 µm and the time series low-pass filtered at 1 Hz with a fourth-order zero-phase Butterworth filter, then converted to a sliding robust z-score (median and median absolute deviation in a 300 s window). The image was tiled with disks of radius ∼4 µm, about the size of one endothelial cell, on a ∼3 µm grid, each tile taking the soft-maximum (log-sum-exp, β = 4) of its pixels. Tile-frames with z ≥ 3.0 were clustered by DBSCAN in (x, y, t) normalized by 3 µm and 1.0 s with minPts 6, and clusters were split at temporal gaps longer than 0.5 s. Events were required to last at least 1.0 s and cover at least 3 tiles, and an automatic raw-brightness gate rejected dim background events. Spatial parameters were fixed in micrometers across sessions. Footprint, duration, peak z and the minimum distance from the footprint to the tracked wall were recorded for each of 35,020 events across 15 sessions.

## Coupling of endothelial calcium events to Plume Events

Triggers were ECCE with onset τ in [−0.5, +5] s of the nearest cue. The near-wall arm comprised every ECCE whose footprint lay within 15 µm of the tracked wall and the far arm every ECCE whose minimum distance from the wall exceeded 25 µm; the intervening annulus was excluded. The response R was the maximum single-trial plume area over [τ, τ + 5] s minus its mean over [τ − 1.5, τ − 0.25] s, on the trigger’s own trial, using the carved-wall detector described above. A per-trigger null took the same cue-relative window on up to 40 neighboring trials (within 20 trials on either side), and a trigger was called coupled if its P ≤ 0.05. Because near-wall and far ECCE differ in amplitude, size and timing, the two arms were balanced within session by propensity-score subclassification^177^ into five quintile strata of the logit propensity on trial index, τ, τ², peak z, log tiles and log duration, with average- treatment-on-the-treated weights applied to the far arm and no near-wall trigger discarded. At least 30 triggers per arm per session were required. The primary statistic was, per mouse, the mean over sessions of the matched near-wall minus far coupled rate, tested by exact one-sided Wilcoxon signed-rank across the five mice. The matcher was calibrated by rerunning it on null triggers, which placed both arms at the nominal rate (4.56% and 4.55%). Specificity was checked with the same matcher on a positive control (ECCE to penetrating-arteriole diameter) and on channels expected to be null (surface ECCE to surface diameter, and a wall translated by 60 µm to plume); the largest positive effect among these sets the residual-bias floor. ECCE amplitude and duration terciles were cut within session and arm, and the cumulative wall-distance ladder (≤ 5, 10, 15 and 25 µm) was pre-specified. Cross-correlation and coherence between the plume signal and ECCE were computed per session on the continuous streams, using the same near-wall (within 15 µm) and far (beyond 25 µm) arms. The cross-correlation was taken on a ±10 s lag grid. Coherence was estimated by multitaper spectral analysis (Chronux^178^; time-bandwidth product 3, 5 tapers, 300 s segments, fpass 0– 0.5 Hz) with per-segment detrending and ordinary-least-squares residualization, and read over 0.1–0.5 Hz. Both were expressed relative to a trial-rotation null, in which the plume stream was realigned against the calcium stream by ±1 to ±20 trials. Both panels are descriptive and are not tested.

## TMN axon imaging

The TMN axon cohort was trained on the same schedule as the plume cohort, four days of Pavlovian followed by eight days of instrumental conditioning, with rhodamine B-dextran injected on Pavlovian day 2 and instrumental days 2 and 8. The axon analyses are aligned to the same task cues. Axon segments were outlined by hand and scored as the mean raw fluorescence of each polygon on every registered frame, without neuropil subtraction. Because these resonant sessions sit close to the detector offset, ΔF/F is understated and not comparable across sessions. The axon trace is therefore expressed as a z- score: the trace relative to a 60 s running median, in robust standard deviations of that residual. The red channel within the same outline was passed through identical arithmetic as a control. Where a session contained several axon segments their mean was used, as they are not independent, and that mean is the session’s unit. An axon event was a rising crossing of z = 3 with a 2 s refractory period, scored by its peak z over the following 2 s. For the comparison of high and low axon activity, trials were sorted by the peak axon z over 0–5 s from the cue and cut into terciles within each session. These analyses are descriptive.

## Analysis of optogenetic stimulation

Train-aligned averages were built per mouse with the blanked lags treated as missing. Clustering used the montage rule above on the baseline-subtracted increase, with a 15 µm band, a 2.5 µm foot ring, a per-frame tracked wall and a per-lag largest-component trim, and the statistic was the peak clustered band area over the response lags. The null was the identical statistic computed on averages anchored on train-free mid-inter-trial windows with a 3 s guard, 1,000–2,000 draws per mouse. P was one-sided and Holm-corrected over the sessions present. A green-excluded test subtracted, per lag, every pixel detected in the structural green channel from the real average and from every draw. Controls comprised a far band at 30–45 µm, a rerun with the first three trains dropped, and dilation-free lags at or after +5.0 s.

## Statistics

The unit of analysis is the mouse, each mouse contributing the mean of its sessions unless stated otherwise; the plume and endothelial calcium cohort is n = 5, the TMN axon cohort n = 4 and the optogenetic cohort n = 8. Per-mouse comparisons used the exact Wilcoxon signed-rank test, whose one-sided floor at n = 5 is 0.03125. Each is reported with the Hodges-Lehmann estimate and the exact signed-rank confidence interval. Where a statistic exists only per mouse, an exact sign-flip test over all 2⁵ sign patterns was used instead. Where trial-level data exist, estimates came from mixed models: fitlme with Satterthwaite degrees of freedom^179^ for continuous outcomes, and fitglme with a logit link and Laplace approximation for binomial outcomes. A random slope by mouse on the tested condition was included wherever that condition varied within a mouse. Singular fits are flagged; the significance marker remains the model’s own P, and a mouse- cluster bootstrap P (see Null models) is reported beside it. Within-session circular-shift and sign-flip tests are reported as secondary, each with its null named. MATLAB functions used were fitlme, fitglme, signrank, radon and iradon. All reported quantifications are mean ± SEM unless otherwise indicated. One test is reported per panel and no correction is applied across panels; Holm correction^180^ within a panel is stated where it is used. Exact P values are given to two significant figures, with *P < 0.05, **P < 0.01 and ***P < 0.001.

## Exclusions

The last cue of each recording was dropped, contaminated trials were excluded from the hit/miss classification, and per-trial analyses used only trials whose analysis window fell inside each recording’s usable diameter record (2,149 of 2,226 trials). Exclusion zones were hand-drawn polygons over bright off-vessel structures and were removed from both the search area and the searchable area used for normalization. ECCE with onsets after the usable record were excluded, and coupling units required at least 30 triggers per arm.

## Software and data availability

Analyses were performed in MATLAB 2025b (MathWorks, Natick, MA) using the Statistics and Machine Learning, Image Processing and Signal Processing toolboxes. Behavior was acquired with custom Bonsai workflows^164^ and a Teensy 4.1, and imaging with Prairie View. Clustering used a C MEX implementation of exact DBSCAN on lattice points whose labels are identical to those of MATLAB’s DBSCAN. Analysis code and the imaging and behavioral data will be deposited in Dryad and made publicly available on publication, with the accession DOI provided at that time (doi:10.5061/dryad.XXXXX).

## Acknowledgements

This work was supported by NIH F32NS134617 (K.L.T.), NIH F31NS143580 (S.J.S.), NIH F99NS125823 (S.F.) and the W. M. Keck Foundation (C.I.M.).

## Author Contributions

K.L.T., S.J.S., S.F., and C.I.M. designed the experiments. K.L.T., S.J.S., S.F., E.T., K.B., C.I.J., and A.N. performed experiments. K.L.T., S.J.S., and S.F. analyzed the data. K.L.T., S.J.S., and C.I.M. wrote the paper.

## Supplementary Figures

**Supplementary Figure 1:**
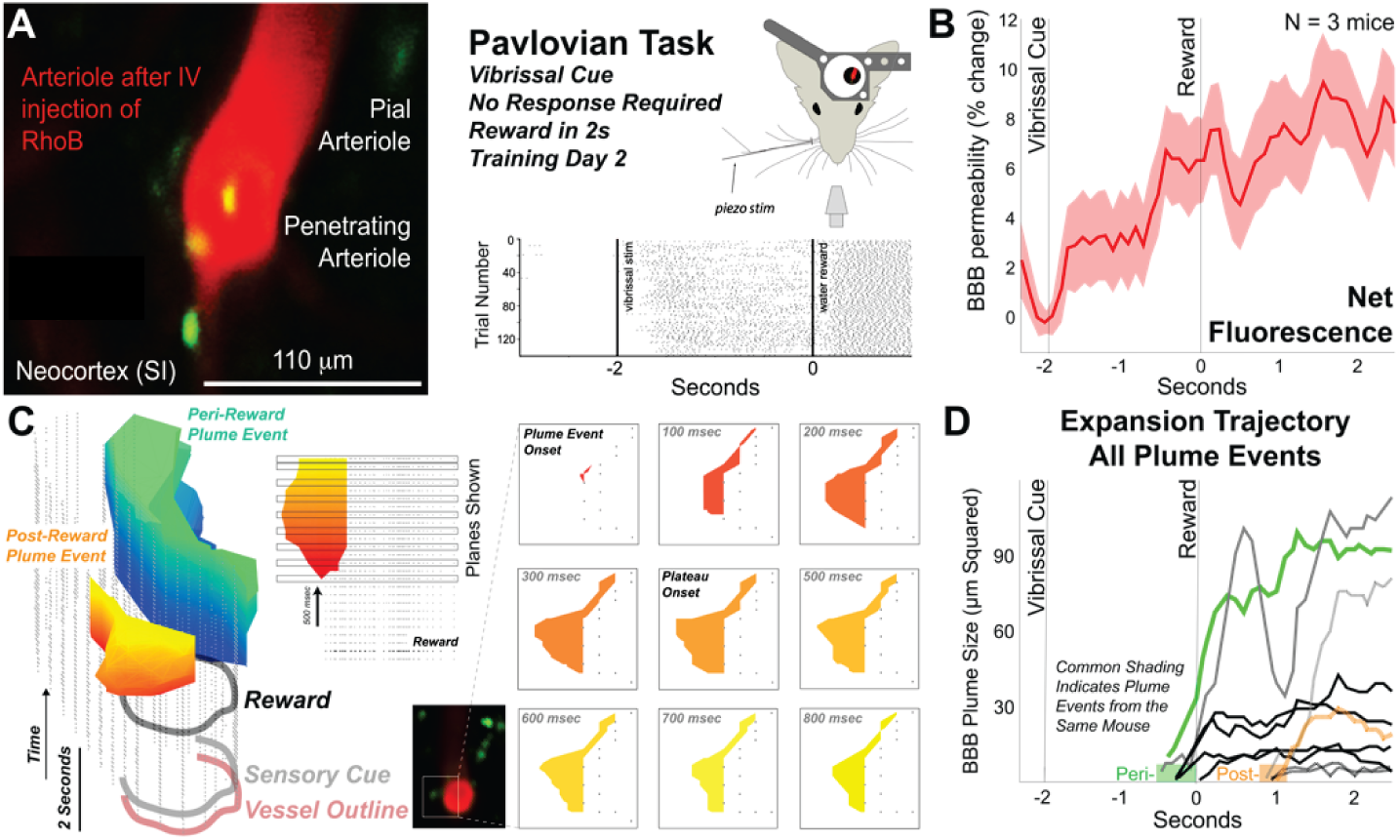
Plume Events are timed to behaviorally relevant epochs. **a**, Two-photon imaging in primary somatosensory neocortex (SI) of a pial-to-penetrating arteriole (PiPA) following tail vein (IV) injection of 70 kDa RhoB-dextran. BBB permeability was imaged on day 2 of training on a Pavlovian variant of a vibrissa cue task. *Raster* shows lick times across trials for one mouse. **b**, BBB permeability measured as change in net fluorescence outside the arteriole during the Pavlovian task (n = 3 mice, 120– 140 trials *per mouse; shading,* 95% confidence intervals). **c**, Plume Events from a Pavlovian task session, rendered as a spatiotemporal volume with a maximal conformation boundary fit (MATLAB). Two distinct Plume Events (*green* and *orange*) are shown with the vessel outline detected in each frame (*points*). Time steps of RhoB expansion into parenchyma are shown for the *orange* Plume Event. **d**, All mice showed Plume Events that onset just before reward (‘pre-reward’) or after (‘post-reward’). The Plume Events shown in *c* are shown as *colored expansion trajectories*.

